# EPHA2 and scavenger receptor-directed trafficking enhances endosomal leakiness and antisense therapy delivery

**DOI:** 10.1101/2025.02.05.635637

**Authors:** Sergi Marco, Peter J. Walsh, Alexey S. Revenko, Peter A. Thomason, Lynn McGarry, A. Robert MacLeod, Sonam Ansell, Dina Tataran, Martin Bushell, Chiara Braconi, Jim C. Norman

## Abstract

The potential for using therapeutic antisense oligonucleotides (ASOs) has been hampered by lack of understanding of how they enter cells and subsequently access their targets. Endocytosis contributes to ASO uptake, but the machinery mediating subsequent ASO trafficking to permit suppression of their target mRNAs has not been described. Here, we show that ASO engagement with a scavenger receptor (CD44) activates the ERK-RSK axis to promote serine phosphorylation of a receptor tyrosine kinase (EPHA2). Serine phosphorylation of EPHA2 permits endocytosis, trafficking, and accumulation of ASOs in nuclear-adjacent endosomes. These endosomes then become leaky, allowing ASOs to escape and effectively suppress target mRNA expression. Inhibition of stress granule-mediated repair of leaky endosomes further enhances ASO effectiveness. These data identify an endocytic route to the nucleus which may be exploited to maximise effectiveness of ASO-mediated therapies.

## Introduction

Antisense oligonucleotides (ASOs) are short single-stranded DNA molecules typically designed to target complementary RNA molecules and regulate their expression. ASO binding to their target RNA forms RNA-DNA hybrid structures, which are recognised by DNA quality control mechanisms of the cell^1-3^. RNase H1, the endonuclease responsible for the degradation of RNA-DNA hybrids^4,5^, then catalyses elimination of the RNA strand, leading to reduction of the encoded protein when ASOs are designed to target mRNA molecules. RNase H1 is mainly localised in the nucleus and mitochondrion, where nascent RNAs are synthesised^6^, but is also present in the cytoplasm where most mature mRNAs are found. Therefore, internalisation and delivery to the appropriate cellular compartment is a key step for effective ASO activity. Various formulations such as encapsulation with liposomes, conjugation with peptides or N-acetylgalactosamine (GalNAc) sugar derivates^2^ have been used to improve the delivery of ASOs and similar molecules. However, these approaches have been reported to evoke pro-inflammatory or other unwanted immune responses^7,8^. The development of phosphorothioate ASOs with constrained ethyl modifications (cET-ASOs) has allowed for the delivery of ASOs by gymnosis (or vehicle-free uptake)^9,10^, which is thought to occur via endocytic routes into the cell.

Following endocytosis, cET-ASOs may be sorted within the endosomal network, and the way in which this occurs dictates their ability to escape, or leak, from endosomal lumens and reach the cytoplasm or nucleus to engage with their target mRNAs and recruit RNase H1^11^. Two obstacles to productive endocytic uptake are post-endocytic return of ASOs to the extracellular milieu or their delivery to degradative compartments (such as lysosomes); both of which reduce opportunities for intact ASOs to leak from endosomes into the cytosolic or nuclear compartments^11,12^. Despite the potential influence of endocytic trafficking on ASO therapeutic efficacy, mechanisms responsible for internalising naked ASOs and the way in which these cargoes are trafficked intracellularly remain largely unclear. A previous study identified and characterised key factors which govern productive uptake of AZD4785; a therapeutic cET-ASO targeting the proto-oncogene *KRAS,* and which is currently under development as a potential anti-cancer therapy^11^. This study highlighted that successful uptake of AZD4785 is associated with the Rab5C endocytic regulator which facilitates ASO trafficking to non-degradative, nuclear-adjacent endosomes. These findings suggest that the positioning of cET-ASOs in endosomes located near the nucleus may enable these molecules to reach their targets.

We have recently described a mechanism in which endosomes are directed to the nuclear vicinity, where they physically associate with the nuclear membrane^13^. This endosome-nuclear interaction, or ‘nuclear-capture’, is mediated by association of nuclear localisation sequences (NLS) in the cytoplasmic tail of the EPHA2 receptor tyrosine kinase (RTK) - a cargo of the captured endosomes – with cytoplasmic components of the nuclear pore. EPHA2 is generally accepted to reside at the plasma membrane to sense contact with neighbouring cells. However, in many tumour types, EPHA2 is overexpressed, leading to accumulation of the receptor on the cell surface^14^, thus leaving plenty spare in its unengaged form to enter the cell via endocytosis to mediate nuclear-capture. Endosomal nuclear-capture is initiated by phosphorylation of Ser^897^ in ligand-unengaged EPHA2. This phosphorylation, along with subsequent endocytosis and nuclear-capture, positions EPHA2 and its downstream signalling activity in nuclear vicinity. Such proximity to the nucleus is a key requirement for EPHA2 signalling to evoke changes in the transcriptional activity of cells undergoing nuclear-capture, further underscoring the importance of positioning in signalling efficacy.

High EPHA2 expression is commonly observed in many tumour types, such as colorectal, ovarian and pancreatic adenocarcinomas (PDAC), which are driven by KRAS activating mutations^15–18^. Mutation of the *KRAS* proto-oncogene is a key driver event in PDAC initiation and, although progressive appearance of loss- and gain-of-function mutations in tumour suppressor genes (such as *TP53*, *CDKN2A* and *SMAD4*) collaborate with mutated *KRAS* to drive tumour aggression^19^, removal of *KRAS* following tumour establishment can provoke PDAC regression in pre-clinical models of the disease^20,21^. Therefore, we investigated the possibility that an endocytic route leading to EPHA2-dependent nuclear-capture of endosomes, which we have previously shown to be active in cells isolated from *KRAS*-driven PDAC^13^, might be exploited to encourage productive uptake of the *KRAS*-targeting cET-ASO (cET-ASO*^Kras^*) in these cells. We describe mechanisms through which EPHA2 collaborates with scavenger receptors to promote internalisation of cET-ASOs and to subsequently direct their trafficking to nuclear-adjacent endosomes which then become leaky, allowing suppression of *KRAS* expression and inhibition of PDAC cell growth.

## Results

### EPHA2 is required for productive uptake of *KRAS*-targeting ASOs

In human PDAC, we found EPHA2 to be strongly expressed in tumour nodules, but largely absent in non-transformed adjacent tissue (Fig. 1A,B; S1A-C). Moreover, indicators of PDAC aggression, including mutation of *KRAS*, *TP53*, *CDKN2A* and *SMAD4*, were associated with increased levels of *EPHA2* (Fig. S1D-H). Consistently, tumours expressing high levels of *EPHA2* displayed diminished overall disease survival and conformed to the most aggressive squamous subtype of the disease (Fig. S1I,J). Importantly, many cells in PDAC tumour nodules exhibited phosphorylation of EPHA2 on Ser^897^ (Fig. 1A,B; S1A,B); an event which is required for nuclear-capture of endosomes^13^. We also examined tumours from the KPC mouse model of PDAC which is driven by pancreas-specific expression of mutants of *Kras* and *Trp53* (*Kras*^G12D/+^; LSL-*Trp53*^R172H/+^; *Pdx1*-Cre) and which recapitulates the aggressive squamous subtype of metastatic PDAC^22^. KPC tumours expressed abundant EPHA2 which was phosphorylated on Ser^897^ thus resembling the human disease in this regard (Fig. 1C, Fig. S1K). To determine the KRAS-dependence of PDAC growth, we allowed cells obtained from KPC tumours to form spheroids and treated them with cET-ASO*^Kras^*(a therapeutic cET-ASO targeting *Kras*). cET-ASO*^Kras^* was taken-up by KPC cells (Fig. S2A) and significantly reduced the volume (Fig. 1D,E) and number (Fig. S2B) of cells in the tumour spheroids they formed, indicating the effectiveness of ASO-mediated inhibition of *Kras* expression to oppose PDAC growth in an appropriate 3D context.

**Figure 1.**
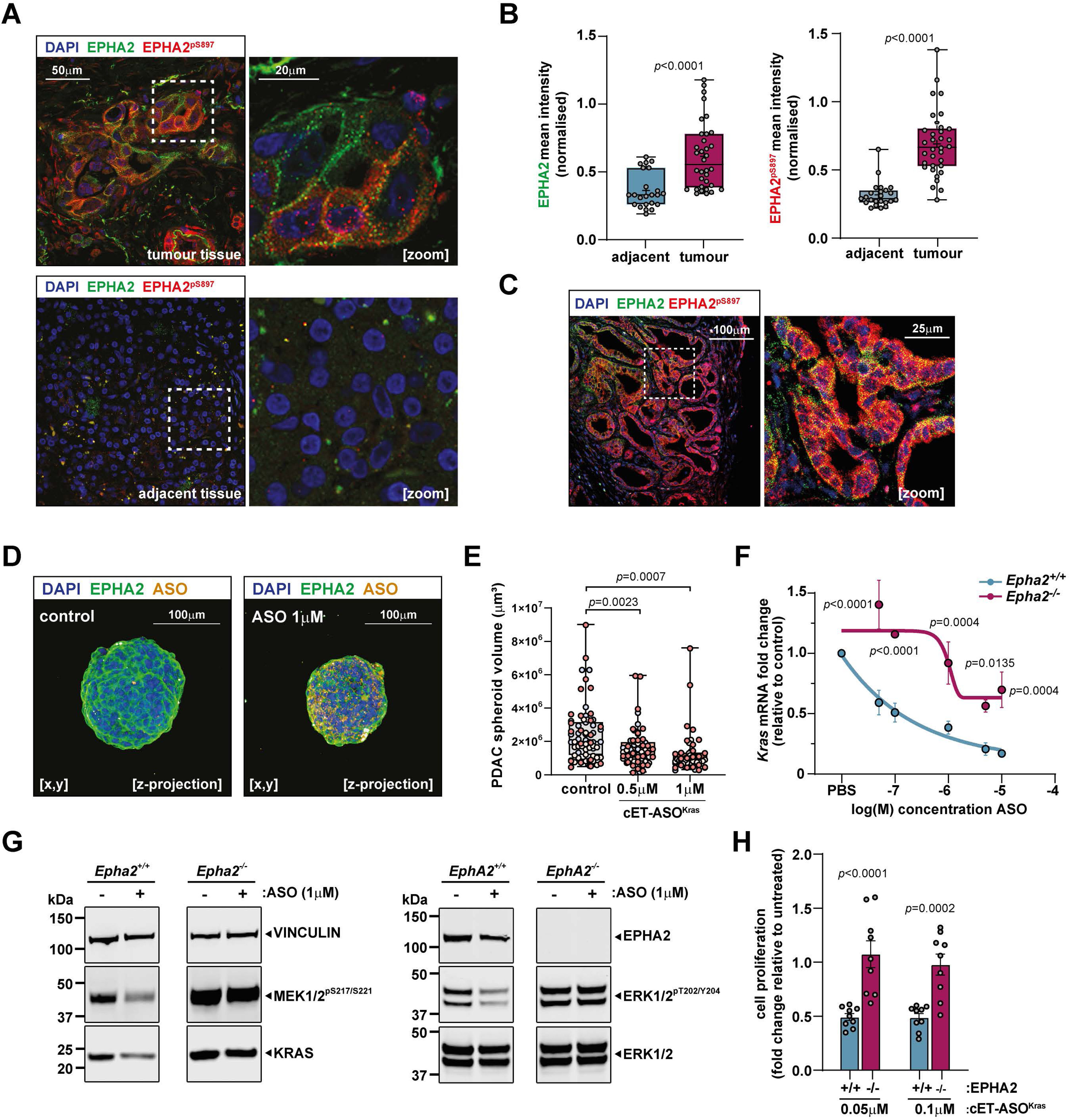
EPHA2 is required for cET-ASO mediated suppression of KRAS in PDAC. (A) PDAC patient tumour tissue immunostaining. Top left and right panels show the staining of EPHA2 (green) and Ser^897^ phosphorylated EPHA2 (red) within the tumour region. Bottom left and right panels show the absence of both EPHA2 and Ser^897^ phosphorylated EPHA2 in non-transformed adjacent tissue in the pancreas. (B) The mean intensity fluorescence value for both EPHA2 (top) and Ser^897^ phosphorylated EPHA2 (bottom) is quantified in both the tumour and adjacent pancreatic tissue across several patient tissue samples. Tumour samples n=6, adjacent non-transformed n=8. Data represented corresponds to five different tumour areas per tumour sample and three different tissue areas per non-transformed sample. Box and whiskers plot, min to max error bars, unpaired t-test (C) Tumour tissue obtained from the KPC (*Kras^G12D/+^*, *LSL-Tp53^172H/+^*, *Pdx1-Cre*) mouse PDAC tumours shows the presence of both EPHA2 and Ser^897^ phosphorylated EPHA2 staining in the tumour region. (D) Confocal imaging Z-projection of immunofluorescence staining of EPHA2 (green) in KPC spheroids treated with cET-ASO^Kras^ (orange) for 72 hours. (E) High-content image analysis of KPC spheroid volume as a proxy for proliferation after treatment with either vehicle or different doses of *Kras*-targeting ASO. Dot colouring indicates each independent experiment. One-way ANOVA, Dunnett multiple comparison test. n=3 (F) Dose response curve of wild-type (*Epha2^+/+^,* blue) or *Epha2* knockout (*Epha2^-/-^*, magenta) KPC cells of *Kras* mRNA expression after 72 hours of treatment with cET-ASO^Kras^. 2-way ANOVA, Sidak multiple comparison test, n=5 individual experiments. (G) Western blot of KPC tumour-derived cell lines from either wild-type (*Epha2^+/+^*) or *Epha2* knockout mice (*Epha2^-/-^*) lysates (left and right respectively) after 72 hours treatment with either vehicle or 1μM cET-ASO^Kras^, showing the changes in the phosphorylation in MEK1/2, ERK1/2and KRAS expression; vinculin used as a loading control. (H) High-content image analysis of either KPC *Epha2^+/+^* (blue) or *Epha2^-/-^* (magenta) cell proliferation after treatment with different doses of *Kras*-targeting ASO for 96 hours. Proliferation index expressed as the fold change of cell number relative to untreated cells. 2-way ANOVA, n=9 individual experiments.

To further investigate involvement of EPHA2 in cET-ASO*^Kras^* uptake into PDAC cells, we used cell lines established from KPC tumours from either wild type (*Epha2^+/+^*) or *Epha2* knockout (*Epha2^-/-^*) mice. We then incubated these cells with cET-ASO*^Kras^* and measured expression of *Kras*. cET-ASO*^Kras^* reduced expression of the mRNA encoding *Kras* in a dose dependent manner; 50% inhibition being achieved at a concentration of approx. 0.1 μM of the cET-ASO (Fig. 1F). In *Epha2* knockout KPC cells, by contrast, the ability of cET-ASO*^Kras^* to suppress *Kras* expression was reduced by more than 100-fold. Western blotting confirmed that 1 μM cET-ASO*^Kras^* suppressed KRAS protein and downstream signalling readouts, including phosphorylation of MEK and ERK, but was unable to do so in *Epha2^-/-^* cells (Fig. 1G). Moreover, addition of a non-targeting ASO (cET-ASO*^NT^*) changed neither *Kras* expression nor its downstream signalling (Fig. S2C,D). Consistently, cET-ASO*^Kras^* suppressed growth of *Epha2^+/+^* cells, but was unable to do so in *Epha2^-/-^* KPC cells (Fig. 1H). The requirement for EPHA2 in ASO efficacy in targeting *Kras* expression was also confirmed by CRISPR-mediated deletion of *Epha2* from both KPC (Fig. S2E-G) and H1299 cells (a human non-small lung carcinoma cell line) (Fig. S2H-J).

### EPHA2-mediated nuclear-capture is required for cET-ASO*^Kras^* to suppress *KRAS* expression

We hypothesised that EPHA2 might contribute to endocytosis of cET-ASO*^Kras^*, and that subsequent EPHA2-mediated nuclear-capture of ASO-containing endosomes may be required for suppression of *Kras* expression. Indeed, cET-ASO*^Kras^* was endocytosed by KPC cells and trafficked to a population of EphA2-positive endosomes located close to and/or in contact with the nucleus (Fig. 2A). Moreover, internalisation of cET-ASO*^Kras^* was significantly reduced in *Epha2* knockout cells, indicating a role for EPHA2 in ASO endocytosis and intracellular trafficking (Fig. 2B). We have previously identified several events necessary for EPHA2-mediated nuclear-capture including: a) phosphorylation of EPHA2 on Ser^897^, b) Rab17-dependent transit of EPHA2-postive endosomes towards the nucleus, and c) engagement of nuclear localisation sequences (NLSs) in EPHA2’s cytotail with the nuclear import machinery^13^. We, therefore, tested the involvement of these processes in the ability of cET-ASO*^Kras^* to suppress *Kras* expression. Firstly, addition of cET-ASO*^Kras^* to KPC cells increased phosphorylation of EPHA2 on Ser^897^ (Fig. 2C). Moreover, the Ser^897^ phospho-defective mutant of EPHA2 (EPHA2^S897A^) displayed significantly reduced ability to rescue cET-ASO*^Kras^*-mediated suppression of *Kras* levels in *Epha2^-/-^* cells (Fig. 2D). Secondly, EPHA2 knockout significantly reduced trafficking of internalised ASO*^Kras^* to RAB17-positive endosomes (Fig. 2E) and, consistently, cET-ASO*^Kras^* -mediated suppression of *Kras* expression and its downstream signalling was compromised in RAB17 knockout cells (Fig. 2F,G). Thirdly, we compared the ability of cET-ASO*^Kras^* to suppress *Kras* expression in EPHA2^-/-^ KPC cells transduced with either wild-type EPHA2 (EPHA2^WT^) or an EPHA2 bearing mutations in its NLS which oppose nuclear-capture of EPHA2-positive endosomes (EPHA2^NLS^)^13^. Although EPHA2^WT^ restored efficient cET-ASO*^Kras^* -mediated suppression of *Kras* levels (Fig. 2H) and MEK/ERK signalling downstream of KRAS (Fig. 2I) to EPHA2 knockout cells, EPHA2^NLS^ was ineffective in this regard. Consistently, cET-ASO*^Kras^* significantly suppressed cell growth in EPHA2^WT^-expressing cells, whereas the proliferation of EPHA2^NLS^-expressing was actually increased following cET-ASO*^Kras^* addition (Fig. 2J). Taken together, these data demonstrate that the ability of cET-ASO*^Kras^* to efficiently suppress expression of its target gene requires EPHA2-dependent endocytosis and subsequent intracellular trafficking to nuclear-captured endosomes.

**Figure 2.**
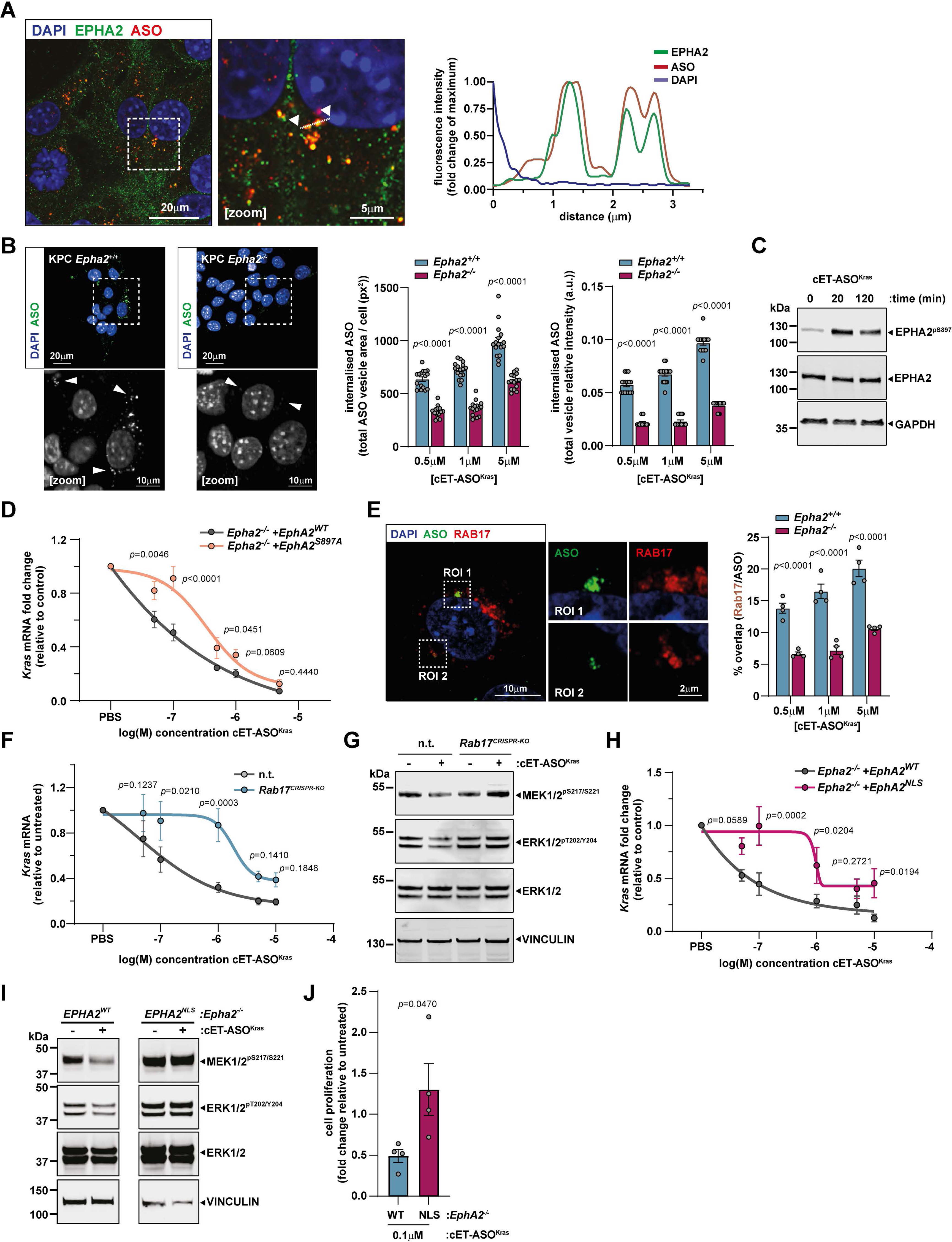
EPHA2-mediated trafficking and nuclear-capture dictates cET-ASOKras efficacy. (A) Immunofluorescence confocal imaging of the internalisation of cET-ASO^Kras^ (red) inside KPC cells after 16 hours of incubation (left panel). Image amplification of the selected area (dotted line frame) showing the colocalisation of cET-ASO^Kras^ and EPHA2 (green) in internalised vesicles (middle panel). Arrowheads and white dotted line indicate the pixels used for the fluorescence intensity histogram profile graph (right panel). (B) Quantification of cET-ASO^Kras^ intracellular vesicle area sum per cell (right panel, left graph) and ASO relative intensity per vesicle (right panel, right graph) in either KPC *Epha2^+/+^* (blue) or *Epha2^-/-^* (magenta) cells treated at different ASO concentrations for 16h. Example micrographs of intracellular cET-ASO^Kras^ vesicles (left panel) in either KPC *Epha2^+/+^* (left) or *Epha2^-/-^*(right) cells. Arrowheads point at internalised vesicles. 2-way ANOVA, n=16 independent samples, each dot corresponds to the average across all the fields of view for each sample. (C) Western blot of KPC *Epha2^+/+^* cells after treatment with 1μM cET-ASO^Kras^ for either 0, 20 or 120 minutes, showing the induction of phosphorylation on EPHA2 Ser^897^ residue. GAPDH was used as a loading control. (D) Dose response curve of *Kras* mRNA expression after 72 hours of treatment with cET-ASO^Kras^ in EPHA2 wild-type rescue (*Epha2^-/-^+ EPHA2^WT^*, grey) or EPHA2^S897A^ mutant rescue (*Epha2^-/-^+ EPHA2^S897^*, salmon) KPC cells. 2-way ANOVA, Sidak multiple comparison test, n=4 individual experiments. (E) Immunofluorescence confocal imaging of the overlap between cET-ASO^Kras^ (green) and RAB17-positive vesicles (red) inside KPC cells after 16 hours of incubation (left panel). Image amplification of the selected area (dotted line frame), and quantification of cET-ASO^Kras^/RAB17 intracellular vesicle overlap in either KPC *Epha2^+/+^* (blue) or *Epha2^-/-^* (magenta) cells treated at different ASO concentrations for 16h (right panel). 2-way ANOVA, n=4 individual experiments. (F) Dose response curve of *Kras* mRNA expression after 72 hours of treatment with cET-ASO^Kras^ in *Epha2^+/+^* non-targeting-CRISPR (grey line), and *Rab17*-CRISPR^KO^ KPC cells (blue line) after 72 hours of cET-ASO^Kras^ treatment. 2-way ANOVA, Sidak multiple comparison test, n=6 individual experiments (G) Western blotting for MAPK signalling (MEK1/2 and ERK1/2) in *Epha2^+/+^* non-targeting-CRISPR and *Rab17*-CRISPR^KO^ KPC cells after 72h of cET-ASO^Kras^ treatment. Vinculin was used as loading control. (H) Dose response curve of *Kras* mRNA expression after 72 hours of treatment with cET-ASO^Kras^ in EPHA2 wild-type rescue (*Epha2^-/-^+ EPHA2^WT^*, grey) or EPHA2 NLS mutant rescue (*Epha2^-/-^+ EPHA2^NLS^*, magenta) KPC cells. 2-way ANOVA, Sidak multiple comparison test, N=7 individual experiments. (I) Western blotting for MAPK signalling (MEK1/2 and ERK1/2) in *Epha2^-/-^+ EPHA2^WT^* and *Epha2^-/-^+ EPHA2^NLS^* KPC cells after 72h of cET-ASO^Kras^ treatment. (J) High-content image analysis of cell proliferation after treatment with cET-ASO^Kras^ for 96 hours in either *Epha2^-/-^+ EPHA2^WT^* (blue) or *Epha2^-/-^+ EPHA2^NLS^* (magenta) KPC cells. Proliferation index expressed as the fold change of cell number relative to untreated cells. 2-way ANOVA, n=4independent experiments.

### Scavenger receptors trigger EPHA2-dependent uptake and trafficking of ASOs

As cET-ASO addition increases phosphorylation of EPHA2 on Ser^897^ (Fig.2C) and because the phospho-defective EPHA2^S897A^ mutant shows a reduced productive uptake of cET-ASO*^Kras^*(Fig. 2D), we interrogated the signalling mechanisms through which cET-ASOs might drive EPHA2 phosphorylation. Several kinases, including AKT and p90RSK can phosphorylate EPHA2’s Ser^897^ downstream of growth factor receptor engagement or cellular stresses such as pro-inflammatory cues^23–26^. Western blotting indicated that KPC cells mounted a rapid (within 15 min) response to cET-ASO (either cET-ASO*^Kras^* or cET-ASO*^NT^*) addition by activating p90RSK, but not Akt (Fig. 3A, Fig. S3A). Furthermore, MAPK signalling (as assessed by phosphorylation of MEK, ERK and JNK, but not p38MAPK), which is established to link surface receptors to p90RSK activation, was also increased within 15 min of cET-ASO addition (Fig. 3A, Fig. S3A,B). Consistently, addition of LHJ685, a potent and selective p90RSK inhibitor (RSKi), completely opposed cET-ASO-driven activation of p90RSK and phosphorylation of EPHA2 at Ser^897^ (Fig. 3B). Finally, p90RSK inhibition significantly decreased the ability of cET-ASO*^Kras^* to suppress *Kras* levels, indicating that phosphorylation of EPHA2 by this kinase is required for productive uptake of the ASO (Fig. 3C). To assess the hierarchy of EPHA2 with respect to the MAPK signalling cascade, we challenged EPHA2 knockout KPC with cET-ASO and found that both ERK, JNK and p90RSK, but not AKT, were activated independently of the cells’ EPHA2 status (Fig. 3D, Fig. S3C). Similarly, CRISPR-mediated deletion of EPHA2 did not reduce cET-ASO-driven phosphorylation of p90RSK or MAPKs in H1299 cells (Fig. S3D). Taken together, these data indicate that cET-ASO-driven activation of MAPK and p90RSK signalling is upstream of Ser^897^ phosphorylation on EPHA2.

**Figure 3.**
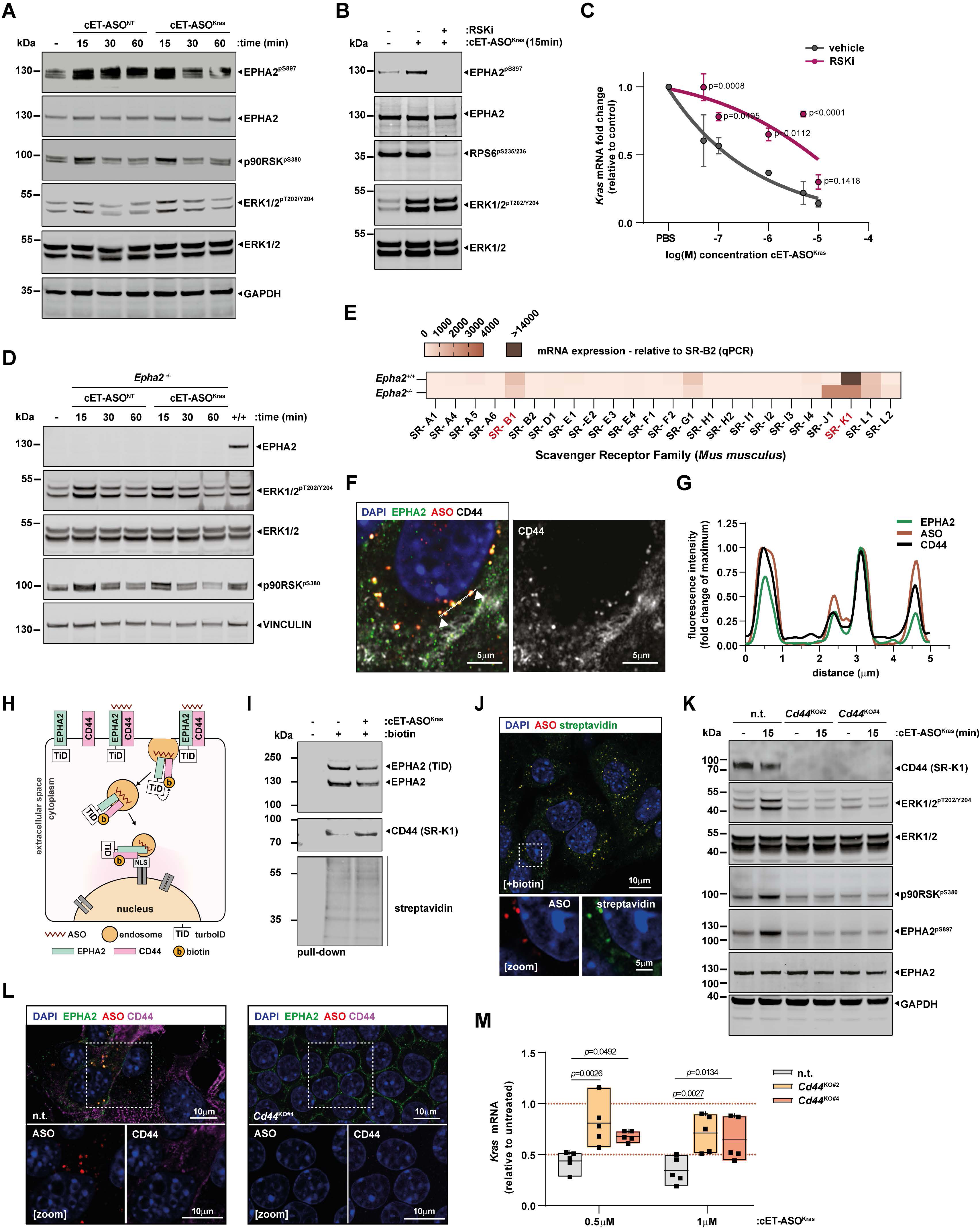
Scavenger receptors trigger EPHA2-dependent uptake and trafficking of ASOs. (A) Western blotting showing induction, by either non-targeting cET-ASO (cET-ASO^NT^) or cET-ASO^Kras^, of the phosphorylation on EPHA2 Ser^897^, p90RSK Ser^380^, and ERK1/2 Tyr^202/204^ in KPC cells. GAPDH was used as a loading control. (B) Western blotting showing the effect of the RSK inhibitor LHJ685 (RSKi) on the cET-ASO^Kras^ -induced phosphorylation of EPHA2 Ser^897^, ERK1/2 Tyr^202/204^ and the ribosomal protein S6 Ser^235/236^ in KPC cells. (C) Dose response curve of *Kras* mRNA expression after 72 hours of treatment with cET-ASO^Kras^ in untreated (grey) or RSK inhibitor treated (LHJ685 10μM, magenta) in KPC cells. 2-way ANOVA, Sidak multiple comparison test, n=3 individual experiments. (D) Western blotting showing the effect of the induction by either cET-ASO^NT^ or cET-ASO^Kras^ of p90RSK Ser^380^ and ERK1/2 Tyr^202/204^ phosphorylations in *Epha2^-/-^* KPC cells and wild-type KPC cells (+/+). Vinculin was used as a loading control. (E) Heatmap showing the levels of mRNA expression (measured by qPCR) of each member of the scavenger receptor family as the fold change of the lowest expressed gene (*Cd36* – SR-B2) in either *Epha2^+/+^* (top) or *Epha2^-/-^* (bottom) KPC cells. Red-labelled names correspond to the highest expressed genes both in KPC and H1299 cells. Outlier values are indicated using dark brown colour. (F) Immunofluorescence confocal imaging of the internalisation of cET-ASO^Kras^ (red), EPHA2 (green) and CD44 (white) in KPC cells after 16 hours of incubation (merge on left panel, Cd44 alone on right panel). Arrowheads and white dotted line indicate the pixels used for the (G) fluorescence intensity histogram profile graph (right panel). (H) Diagram depicting the mechanism of EPHA2-TurboID fusion protein (TiD) labelling of proximity proteins including CD44 after cET-ASO^Kras^ addition. (I) Western blotting showing the pulldown of EPHA2-TurboID biotin-labelled proteins corresponding to KPC cells that were untreated or incubated with biotin (100μM) and/or cET-ASO^Kras^ for 16 hours. Western blots show the pulldown of EPHA2-TurboID, wild-type EPHA2, and CD44. Streptavidin detection of biotinylated proteins is used as a control for gel loading. (J) Immunofluorescence confocal imaging of cET-ASO^Kras^ internalised vesicles (red) and biotin-labelled proteins (green) in EPHA2-TiD expressing cells. (K) Western blotting showing the effect of *Cd44* CRISPR knockout KPC cell lines on the cET-ASO^Kras^-induced activation of ERK1/2 Tyr^202/204^, p90RSK Ser^380^ and EPHA2 Ser^897^ phosphorylation. GAPDH was used as loading control. (L) Immunofluorescence imaging of the overlap of intracellular CD44 and internalised ASO in control and *Cd44^KO#2^* KPC cells. (M) *Kras* mRNA expression in either non-targeting or *Cd44^CRISPR-KO^* KPC cell lines treated with either 0.5μM or 1μM cET-ASO^Kras^ for 72h. 2-way ANOVA, Dunnett multiple comparison test, n=5 individual experiments.

We next considered whether a receptor-mediated mechanism might be responsible for cET-ASO-driven activation of MAPK/p90Rsk signalling and the resulting EPHA2 phosphorylation. Scavenger receptors (SRs) are a family of cell surface proteins established to mediate antigen uptake by myeloid cells, as well as blood lipid clearance by hepatocytes in the liver^27,28^. More recently SRs have been shown to participate in internalisation of polyanionic molecules (such as ASOs)^29,30^ and, in some cases, to initiate MAPK signalling^31,32^. We, therefore, screened for SR expression in both KPC (Fig. 3E) and H1299 (Fig. S3E) cells, which highlighted *SCARB1* (SR-B1) and *CD44* (SR-K1) as the most highly expressed members of this receptor family in both cancer cell types. Furthermore, both *CD44* and *SCARB1* were detected at and nearby the plasma membrane of cells in the tumour compartment of human PDAC (Fig. S3F,G), and expression of *CD44*, like *EPHA2*, is highest in the most aggressive squamous PDAC subtype^33^ (Fig. S3H).

In KPC cells, cET-ASO*^Kras^* was endocytosed and trafficked to endosomes that were positive for both EPHA2 and CD44 and which congregated in the vicinity of the nucleus (Fig. 3F,G). We then considered whether engagement with cET-ASOs may contribute to bringing the RTK and SR into physical proximity. To test this, we use a BioID approach in which we tagged the C-terminal domain of EPHA2 with a TurboID ligase which, upon provision of a biotin substrate, catalyses biotinylation of nearby (within 10nm) proteins (Fig. 3H). Addition of cET-ASO*^Kras^* to KPC cells increased the ability of a EPHA2-turboID fusion protein to biotinylate CD44 (Fig. 3I), and biotin-labelled proteins could be seen in intracellular vesicles (Fig. 3J) following ASO addition, indicating that cET-ASOs triggers increased physical association of EPHA2 and SRs which is associated with their endocytosis.

These associations prompted us to investigate the role played by SRs in key events including cET-ASO internalisation, the MAPK/p90RSK signalling downstream of cET-ASO addition, and the ability of cET-ASO*^Kras^* to suppress *KRAS* expression. Firstly, CRISPR-mediated deletion of either *Cd44* or *Scarb1* in KPC cells (Fig. S4A,B) opposed cET-ASO-driven activation of MAPKs, p90R and phosphorylation of EPHA2 at Ser^897^ (Fig. 3K, Fig. S4C). Secondly, in cells lacking CD44 we were unable to detect vesicles containing either ASOs, internalised EPHA2 or both these molecules (Fig. 3L). Finally, CRISPR-mediated deletion of *Cd44* or *Scarb1* opposed the ability of cET-ASO*^Kras^* to suppress *Kras* expression in KPC (Fig. 3M, Fig. S4D) and H1299 cells (Fig. S4E-G). Taken together, these data indicate that cET-ASOs activate SRs leading to p90RSK activation, physical association of CD44 and EPHA2 and phosphorylation of EPHA2 at Ser^897^. This permits internalisation and co-trafficking cET-ASO, an SR and phosphorylated EPHA2-pSer^897^ to juxta-nuclear endosomes which, in turn, enables productive uptake of cET-ASO*^Kras^* and effective suppression of its target mRNA.

### EPHA2 is required for cET-ASO trafficking to leaky endosomes

Escape, or leakage, of endocytosed ASOs across endosomal membranes is thought to facilitate their movement to the cytoplasm or nucleus, thus permitting interaction with their target mRNAs. However, while acquisition of endosomal leakiness favours ASO release, it also triggers recruitment of membrane repair effectors including ESCRT components and/or galectins (Galectin-3, −8 and −9)^34–39^. More recently, endo-lysosomal damage has been shown to trigger assembly of ribonucleoprotein granules (RNP), stress granules (SG)^40^ - liquid-like membrane-less condensates which assemble in the cytosol as a response to cell stressors^41^. G3BP1, a core SG component, is recruited to leaky endosomes and contributes to plugging and repair of the damaged membranes. We, therefore, deployed galectin-9 and G3BP1 as probes to identify leaky membranes and, thus, monitor potential sites of endosomal escape of internalised cET-ASOs. First, addition of cET-ASOs markedly increased the size of both galectin-9 and ASO positive intracellular vesicles, as well as the overlap between them in KPC *Epha2^+/+^* cells, but not *Epha2^-/-^* cells (Fig. 4A,B; Fig. S5A,B). Indeed, following cET-ASO addition many intracellular structures were triply positive for cET-ASO, galectin-9 and G3BP1 and predominantly located at, or nearby, the surface of the nucleus (Fig. 4C). Recruitment of G3BP1 to these endosomes was again EPHA2-dependent, as most KPC *Epha2^-/-^* cells failed to generate any positive G3BP1 cytoplasmic condensates (Fig. 4D,E) and, when present, these were reduced both in number and size (Fig. 4F, Fig. S5C-F). Moreover, both EPHA2 and EPHA2^pS897^ could be observed to overlap with G3BP1- and galectin-9 positive intracellular structures (Fig. S5G,H), suggesting their association with the nuclear-capture endosomal pathway. Taken together, these data indicate that cET-ASOs, via activation of SRs and their downstream signalling, promote EPHA2-pSer^897^-dependent endocytosis and trafficking of cET-ASOs which culminates at nuclear-proximal/nuclear-captured compartments. These compartments then become leaky to permit endosomal escape and productive uptake of cET-ASO*^Kras^*.

**Figure 4.**
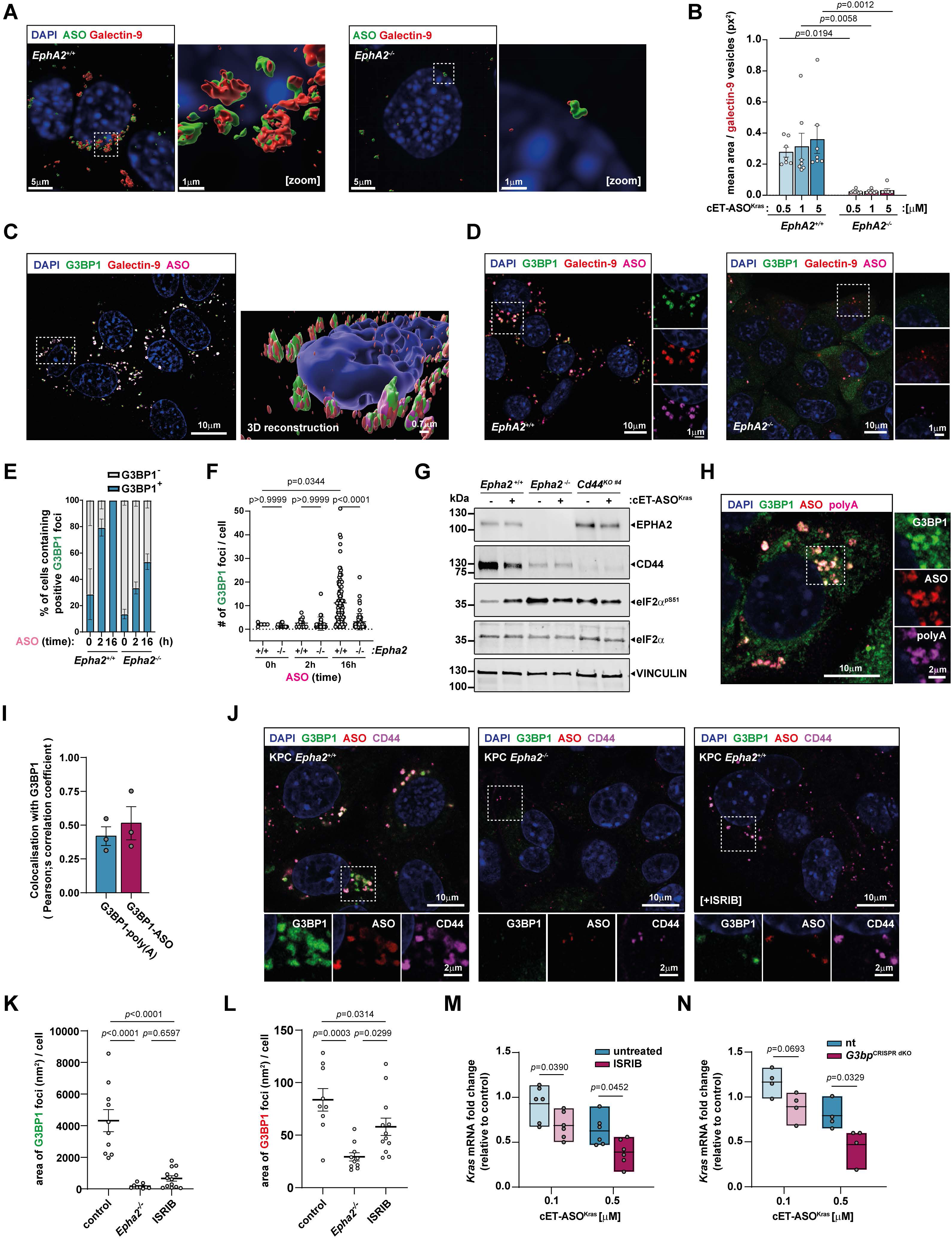
EPHA2 is required for trafficking of cET-ASOs to leaky endosomes. (A) 3D reconstruction images of Galectin-9 (red) overlap with internalised cET-ASO^Kras^ (green) vesicles in KPC cells. Image amplifications correspond to the white dotted line frames. (B) Quantification of the overlap between Galectin-9 and ASO intracellular structures in either *Epha2^+/+^*(blue) or *Epha2^-/-^* (magenta) KPC cells, after treatment with either 0.5, 1 or 5 μM concentrations of cET-ASO^Kras^ for 16 hours. One-way ANOVA, Tukey multiple comparison test, n=7. (C) 3D reconstruction images of Galectin-9 (red) overlap with internalised cET-ASO^Kras^ (magenta) and G3BP1-GFP (green) in KPC cells. Image amplifications correspond to the white dotted line frames. (D) 3D reconstruction images of galectin-9 (red) overlap with internalised cET-ASO^Kras^ (magenta) and G3BP1-GFP (green) in either *Epha2^+/+^* or *Epha2^-/-^* KPC cells. Image amplifications correspond to the white dotted line frames. (E) Percentage of cells that contain at least one G3BP1 positive condensate in either *Epha2^+/+^* or *Epha2^-/-^* KPC cells after treatment with cET-ASO^Kras^ for 0, 2 or 16 hours. (F) Average number of G3BP1-positive structures per cell, in either *Epha2^+/+^* or *Epha2^-/-^* KPC cells after incubation with ASO cET-ASO^Kras^ for 0, 2 or 16 hours. 2-way ANOVA, Tukey multiple comparison test, Dots correspond to the different fields of view obtained in three individual experiments. One-way Anova – Tukey test. (G) Western blotting of the effect on eIF2a phosphorylation on Ser^51^ by cET-ASO^Kras^ in either *Epha2^+/+^* or *Epha2^-/-^* KPC cells after incubation with ASO cET-ASO^Kras^ for 4 hours. Vinculin was used as loading control. (H) Micrograph showing fluorescence *in situ* hybridisation with a poly-d(T) probe (magenta) and its overlap with G3BP1-GFP (green) and cET-ASO^Kras^ (red) after 16h treatment in KPC cells. (I) Quantification of the overlap using Pearson’s correlation coefficient between the G3BP1-GFP and mRNAs (poly-d(T)) staining, and G3BP1-GFP and cET-ASO^Kras^ staining. n=3 individual experiments. (J) Micrographs of immunofluorescence confocal imaging showing the overlap between G3BP1-GFP, internalised cET-ASO^Kras^ and intracellular CD44 in either *Epha2^+/+^*, *Epha2^-/-^* or ISRIB (1μM) treated *Epha2^+/+^* KPC cells. (K) Quantification of the average sum of the area of intracellular G3BP1-GFP condensates and (L) ASO vesicles per cell, in cET-ASO^Kras^-positive vesicle cells. Dots correspond to the different fields of view obtained in three individual experiments. One-way Anova – Tukey test.. (M) Effect of cET-ASO^Kras^ in *Kras* mRNA expression as fold change of control in untreated or ISRIB-treated (1μM) KPC cells. 2-way ANOVA, Sidak multiple comparison test, n=6 individual experiments. (N) Effect of cET-ASO^Kras^ on *Kras* mRNA expression as fold change of control in either non-targeting (n.t.) or double *G3bp1/G3bp2* CRISPR knockout (G3BP^dKO^) KPC cells for 72h. 2-way ANOVA, Sidak multiple comparison test, n=4 individual experiments.

### EPHA2-dependent uptake of cET-ASO*^Kras^* is enhanced by inhibition of endosomal repair

A range of cellular stresses, including viral infection and nutrient, ER and oxidative stresses activate kinases - such as PKR, PERK, GCN2 and HRI - that increase phosphorylation of eIF2α which, in turn, triggers SG assembly^42^. Addition of cET-ASO increased phospho-eIF2α in KPC cells (Fig. 4G); a response that did not occur in *Epha2-* or *Cd44*-knockout cells, indicating that EPHA2-dependent internalisation of cET-ASOs and their subsequent delivery to nuclear-captured endosomes constitutes a stress with the potential to drive SG assembly. Fluorescence *in situ* hybridisation with a poly-d(T) probe indicated very close colocalisation of polyadenylated mRNAs and G3BP1 with internalised cET-ASOs (Fig. 4H,I). Similarly, after cET-ASO addition, G3BP1/galectin-9-positive structures were found associated with the SG marker, eIF3b (Fig. S5I)^43^. Thus, the cET-ASO-positive endosomes which accumulate in EPHA2-expressing cells following cET-ASO addition appear to be in very close association with *bone fide* SG markers.

In addition to their role in regulating mRNA translation under stress conditions, SGs have recently been shown to contribute to plugging holes in endo-lysosomal membranes which have been damaged by internalised mycobacteria^40,44^. Interestingly, by repairing membrane damage, SGs can reduce leakage of mycobacteria from the endosomal system to restrict their pathogenicity. We proposed that formation of SGs at endosomal damage sites may similarly restrict cET-ASO efficacy. ISRIB (integrated stress response inhibitor) – a compound established to oppose SG assembly by blocking the consequences of eIF2α phosphorylation^45,46^ – was used to test this hypothesis. As expected, addition of ISRIB opposed cET-ASO-driven SG assembly and recruitment of G3BP1, maintaining a significant uptake rate of cET-ASOs into endosomes (Fig. 4J-L; Fig. S5J,K). Furthermore, as well as inhibiting SG formation, ISRIB significantly increased the ability of cET-ASO*^Kras^* to suppress expression of *Kras* mRNA (Fig. 4M).

SG assembly may also be opposed by combined knock out of its core components *G3BP1* and *G3BP2^41^*. CRISPR deletion of both genes (*G3bp*^CRISPR^ ^dKO^) (Fig. S5L), strongly enhanced the ability of cET-ASO*^Kras^* to suppress *Kras* expression (Fig. 4N) in KPC cells. Taken together, these data indicate that EPHA2-dependent trafficking of cET-ASO*^Kras^* to leaky endosomes in the nuclear proximal region is necessary for the ASO to efficiently suppress its target gene, and that ASO efficiency may be further enhanced by reducing SG-mediated endosomal repair mechanisms.

## Discussion

In this study we have identified a pathway transporting a cET-ASO with therapeutic potential to leaky endosomes. This endocytic journey is initiated by cET-ASO engagement with a scavenger receptor (such as CD44) which activates p90RSK to phosphorylate a receptor tyrosine kinase, EphA2. Phosphorylation of EphA2 at Ser^897^, in turn, permits endocytosis and Rab17-dependent centripetal co-transport of endosomes containing cET-ASO, CD44 and EPHA2. An NLS in EPHA2’s cytotail then enables nuclear-capture of these endosomes which become leaky, allowing the cET-ASO to escape from the endosomal lumen and effect suppression of its *Kras* target mRNA. Importantly, these cET-ASO-driven signalling and trafficking events promote recruitment of mRNA stress granule components to nuclear-captured cET-ASO/EPHA2/CD44-containing endosomes. Stress granules have a newly-established role in plugging damaged endosomal membranes and, consistently, inhibition of stress granule assembly opposes endosomal repair and further enhances cET-ASO uptake and suppression of its target mRNA. These events are summarised in the scheme presented in figure 5.

**Figure 5.**
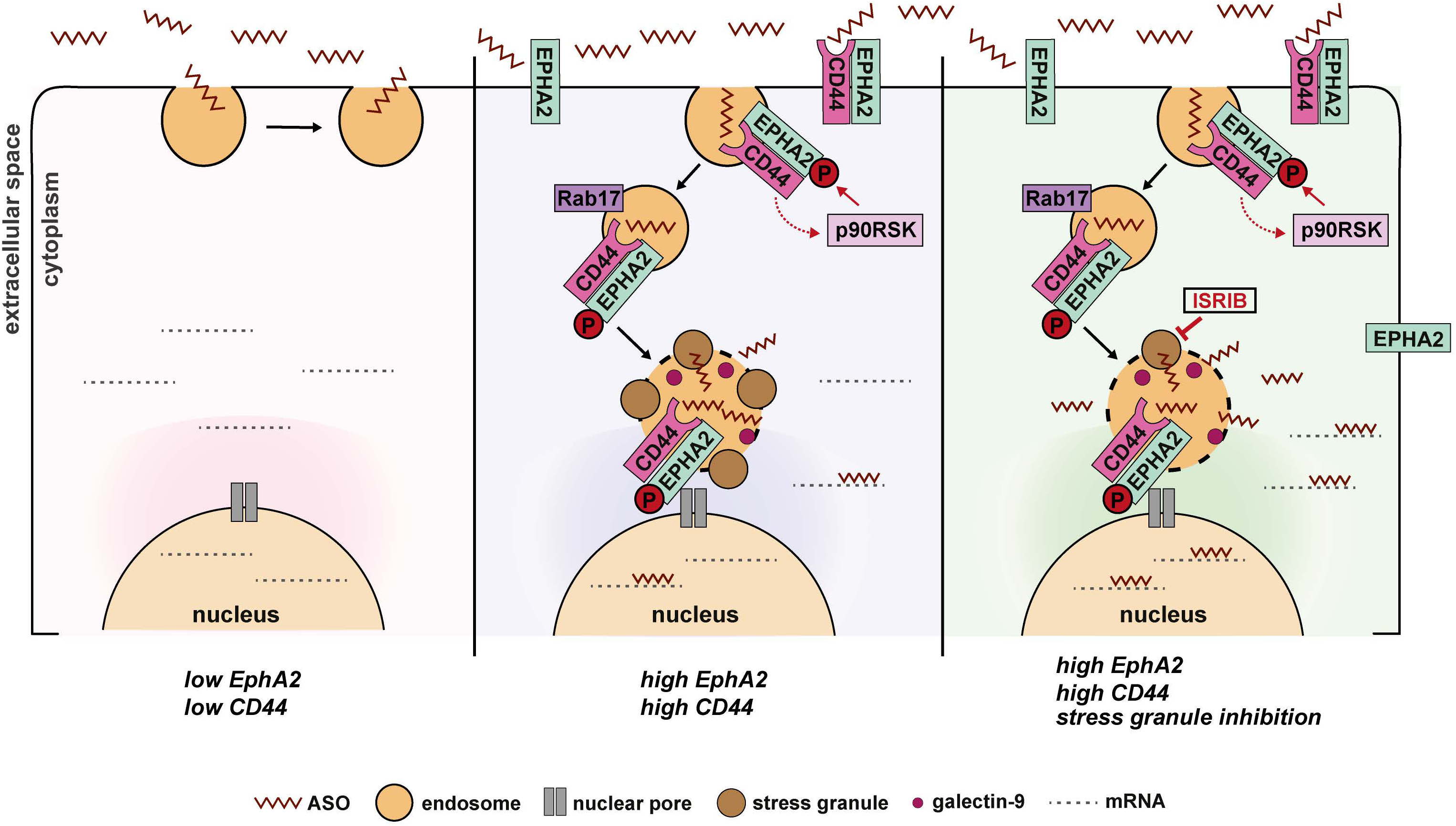
(A) Schematic diagram showing the ASO uptake and mRNA-targeting efficacy in cells expressing low levels of EPHA2 and CD44 (left), in cells expressing high levels of EPHA2 and CD44 (middle), and in cells with high levels of EPHA2 and CD44 expression combined with suppression of stress granule formation using the ISR inhibitor ISRIB.

SR-dependent activation of a signalling cascade driving EPHA2 phosphorylation precedes cellular cET-ASO accumulation, indicating that ligand-engagement of an SR and the consequent activation of p90RSK is an initiating event in cET-ASO endocytosis and trafficking. As their name implies, members of the SR family bind to and participate in the endocytosis of a wide range of endogenous and xenobiotic molecules, which often involves downstream activation of MAPKs^31,47,48^. This process is most well-understood in professional antigen presenting cells (dendritic cells and macrophages) where SRs mediate antigen uptake and trafficking for presentation on MHC class II^49^. In dendritic cells, SRs also contribute to cross-presentation of exogenous (endocytosed/phagocytosed) antigens via a ‘cytosolic pathway’ where antigens are processed by the proteasome for presentation on MHC class I^50–52^. How these antigens reach the cytosol is a matter of some debate. However, it is now apparent that endosomal leakage/escape of endocytosed/phagocytosed antigens can allow them to access the cytosolic proteasome complex^35^. Thus, antigen cross-presentation now provides *de facto* evidence for a physiological process linking SR-mediated internalisation and trafficking to leaky endosomes. However, it is unlikely that SRs are present on PDAC cells to present antigen, and it is more likely that other functions, such as maintenance of cancer cell stemness or uptake of nutrients (such as cholesterol and fatty acids), are likely responsible for providing growth advantage to cancer cells ^53^. Thus, we propose that a receptor which has been selected to be abundantly expressed in tumours is also a gatekeeper to an endocytic pathway which may be exploited to deliver therapeutic molecules to aggressive tumours.

RTKs, such as EGFR1, have previously been shown to mediate cET-ASO endocytosis^54^. However, it is not only the uptake of cET-ASOs, but the way they are subsequently trafficked that dictates whether they can reach their target mRNAs.

Indeed, our evidence highlights a role for a particular RTK, EPHA2, in both uptake and, importantly, the subsequent trafficking of a cET-ASO to leaky endosomes. EPHA2 is a multifunctional protein with established roles both as a mediator of repulsive cues during tissue patterning and as a driver of invasion and metastasis in cancer^55^. So why does this RTK play such a pivotal role in engendering a substantial population of leaky endosomes? EPHA2’s signalling and trafficking differ markedly between its contribution to tissue patterning during development and its pathological role driving cancer cell invasiveness. Very high expression levels of EPHA2 lead to a population of ligand-unengaged receptor which can become phosphorylated on Ser^897^ in response to certain stimuli, such as activation of the cMET RTK^13,56^. Most pertinently, in cancer, genomic data indicate that tumours carrying KRAS-activating mutations (the overwhelming majority of PDAC) have particularly high expression of EPHA2 (Fig. S1D), positioning such tumours as ideal candidates for cET-ASO therapies generally. Moreover, because mutation of *KRAS* is an early event in PDAC carcinogenesis and is apparent in pre-neoplastic PanIN lesions^57^, an assessment of the EPHA2-Ser^897^ phosphorylation status of PanINs will be important to evaluate the potential of cET-ASOs as a preventative strategy for PDAC.

Unlike the tyrosine autophosphorylation evoked by engagement of EPHA2 with its ephrin ligands, phosphorylation of EPHA2’s Ser^897^ which can occur in response to activation of cMET (but also CD44 as we describe here) does not terminate signalling but allows assembly of a signalosome comprising Ephexin4 and RhoG^58^. This cascade activates Rac1 to drive actin polymerisation and invasive cell migration. Moreover, EPHA2’s drive to invasive cell migration is enabled by nuclear-capture of the endosomes upon which the EPHA2/RhoG signalosome is assembled^13^. This is because nuclear-capture localises signalling downstream of RhoG to the immediate vicinity of the nucleus, thus driving activation of the MRTF/SRF transcription factors to support and maintain invasive phenotypes. The nuclear proximity of EPHA2/cET- ASO-containing leaky endosomes, and the role played by EPHA2’s NLSs in this, mechanistically links nuclear-capture to acquisition of endosomal leakiness.

But how could nuclear-capture influence endosomal leakiness? Endosomal leakiness varies between compartments. Indeed, compartments with early endocytic/recycling characteristics being reported to have the highest probability for cargo escape^59^. Consistently, manipulations leading to cargo accumulation in early endosomal/recycling compartments have also been shown to increase the activity of ASOs^60^. Our data indicate that the compartment in which cET-ASOs accumulate in an EPHA2-dependent manner, in addition to being RAB17-positive (Fig. 2E) stain for several other recycling endosome markers, including RAB11, 14, and RCP (Fig. S6). This observation is consistent with nuclear-captured endosomes possessing early endocytic/recycling compartment and, therefore, leaky characteristics. Therefore, we propose that the activation of EPHA2 non-canonical signalling favours the trafficking of cET-ASO into recycling-type endosomes, a population with a propensity to become leaky. Moreover, it is possible that tethering these leaky endosomes to the nuclear surface may concentrate cET-ASOs in the immediate vicinity of the nucleus, thus maximising their deliver to the main location of RNaseH1 and nascent mRNAs.

Chemical manipulations of the endosome trafficking and repair machinery that have been shown to enhance oligonucleotide delivery have been limited by toxicity^61^, whilst screening approaches to identify new components of the endocytic machinery that might enhance antisense oligonucleotide activity have not been investigated in detail. However, our study introduces another actor, the stress granule, which appears to make important contributions to productive uptake of an cET-ASO. SGs, although well-established as mRNA translation regulatory organelles ^62,63^, are now being proposed to participate in endomembrane repair^40,44^, and therefore may have a role regulating endosomal leakiness. Interestingly, the upstream signalling that shuts down translation following nutrient and proteotoxic stresses and which drives SG assembly is well-characterised and, therefore, amenable to pharmacological intervention. Indeed, our observation that addition of ISRIB to oppose SG assembly and increase cET-ASO efficacy highlights the potential of the exploring the pharmacological tools established to manipulate the mRNA translation machinery for enhancing antisense oligonucleotide-based therapies. Furthermore, we anticipate that further studies focused on elucidating the emerging mechanistic relationship between endocytosis, mRNA translation and endomembrane repair in normal and pathophysiologic situations will provide valuable insight into how to deliver therapeutics in a targeted and effective way.

## Methods

### Tissue immunofluorescence

Immunofluorescence staining was performed on paraffin-embedded tissue sections of patient pancreatic adenocarcinoma (PDAC) tumours or KPC mouse PDAC tumour sections. Human tissues were retrieved through patients undergoing surgery with curative intent within Greater Glasgow and Clyde NHS hospitals through the Glasgow Biorepository (N.526) (n=7). KPC tissue samples were obtained from archived paraffin blocks used in previous studies^56^.

Briefly, paraffin was removed from tumour section containing slides using sequential steps with xylene, 100%, 95%, 80% ethanol washes and a final step with H2O rehydration. Antigen retrieval was performed in citrate buffer pH6.0, and samples were placed in a water bath at 100°C for 20 min, and allowed to cool down at RT for a further 20min. Slides were washed three times with TBS and samples were incubated with blocking buffer (1% normal goat serum, 0.1% Triton X-100 in TBS) for 1h at RT. Primary antibodies were incubated ON at 4°C. Next day, slides were washed 3 times with TBS and the corresponding secondary antibodies together with DAPI were incubated on each slide for 1h at RT. Next, samples were washed 3 times with TBS buffer and coverslips were mounted afterwards using Fluoromount G mounting medium. Samples were visualised using a Zeiss LSM880 microscope with a 20x/0.8NA objective and Zen Black 2.3 software, sequential excitation was used, pixel size was 0.06 µm. Image analysis to measure the mean intensity of each fluorophore was performed across the different patient tissue in different fields of view for each patient using Fiji-ImageJ software (2.6.0).

### Cell culture

KPC and H1299 cells were cultured in DMEM, high glucose, GlutaMAX™, supplemented with 10% of FBS (v/v). Cells were cultured at 37°C in a controlled humidified environment containing 5% CO2, and cultures were split every 3–4 day periods using trypsin/EDTA solution. Cells were tested for the presence of mycoplasma and H1299 cells were validated using Promega GenePrint 10 system (STR multiplex assay) at the Molecular Technology Services (CRUK-SI).

### KPC PDAC spheroids

Agarose 9×9 micro-molds were prepared as per the manufacturer’s instructions (MicroTissues® 3D Petri Dish® micro-mold, Merck) and placed in each well of a 12-well cell culture plate. KPC cells were trypsinised and 4.5×10^5^ cells were seeded into each agarose micro-mold using 190ul of medium. Cells were allowed to enter the micro-wells for 2h at 37°C and the medium was topped-up immediately after covering the whole micro-mold. Cells were allowed to form spheroids for three days and on DIV3, cET-ASO^Kras^ was added to each corresponding well. On DIV7 cells, the spheroids were fixed while inside the micro-molds using 4% paraformaldehyde for 1h at RT. Next, the spheroids were popped out from the microwells using PBS, collected and transferred to an empty 96-well plate for manipulation. Spheroids were then incubated with blocking buffer (0.1% Triton X-100, 2% BSA, PBS) for 30min at RT. Primary antibodies were diluted in blocking buffer and incubated with the spheroids ON at 4°C. Next day, three 30-minute washes with blocking buffer were performed for each set of spheroids and then samples were incubated with the corresponding secondary antibodies as well as DAPI. Samples were incubated ON at 4°C. Next day, the spheroids were washed 3 times for a 30-minute period each time and finally samples were mounted with FUnGI solution (60% glycerol, 2.5M fructose, H2O) on a slide, using a frame seal (Bio-Rad) and a coverslip. Samples were visualised using either a Zeiss LSM880 confocal microscope as above, or using Revvity OperaPhenix automated confocal microscope. Z-stacks using a 1 µm step were obtained for each spheroid and used to measure spheroid cell number and spheroid total volume. Data shown corresponds to the individual spheroids pooled from independent experiments. Image analysis was performed using Harmony High-Content image analysis software (Revvity).

### cET-ASO dose-response curve

KPC or H1299 cells were seeded at a density of 1.5×10^5^ cells for each condition in a 6-cm plate. Cells were allowed to attach and proliferate overnight. Next day, cells were treated with either vehicle (PBS) or cET-ASOs at 0.05, 0.1, 1, 5 and 10μM concentrations for 72h. Next, RNA was harvested using Trizol (ThermoFisher). 1μg of RNA was used for RT-PCR to produce cDNA (High-capacity RNA to cDNA kit, ThermoFisher) following the manufacturer’s recommended protocol. *Kras* mRNA expression was assessed using quantitative PCR (qPCR) using QuantiNova SYBR Green RT-PCR Kit (Qiagen), a CFX1000 thermocycler (BioRad) and CFX Maestro software (BioRad). RPLP0 ribosomal RNA was used as reference gene. *Kras* mRNA expression is expressed relative to untreated cells for each condition across independent experiments.

### Western blotting

KPC or H1299 cells were seeded at a density of 3×10^5^ cells in a 6-cm plate for long experiments (72h) or 1×10^6^ cells for short experiments (15min – 16h). Cells were treated with either vehicle (PBS) or cET-ASOs accordingly and protein was harvested using RIPA buffer supplemented with protease and phosphatase inhibitors (Halt™ Protease and Phosphatase Inhibitor Cocktail, ThermoFisher) at endpoint. Protein lysates were sonicated and insoluble fractions were discarded after scentrifugation at 10000rpm at 4°C for 5min. Protein concentration was measured using a BCA assay (ThermoFisher). Uniformly concentrated samples were separated by electrophoresis, using NuPAGE 4-12% Bis-Tris gels (ThermoFisher) and proteins were transferred to a nitrocellulose membrane using the Trans-Turbo transfer system (BioRad). Membranes were further incubated with blocking buffer (5% BSA in TBS 0.1% Tween20) for 1 hour at RT. Corresponding antibodies were incubated ON at 4°C on a rocker. Membranes were washed the following day three times with TBS-T at RT. The corresponding secondary antibodies were incubated in blocking buffer for 1 hour at RT. Three washes with TBS-T were subsequently performed and protein expression was evaluated either using the Odyssey CLx Imager (Li-Cor) or using an ECL substrate (Supersignal West Femto substrate, ThermoFisher) and imaging with Chemidoc imaging system (BioRad).

### Cell proliferation assay

KPC cells were seeded in 96-well optical bottom plates using 750 cells per well. Cells were allowed to attach and proliferate overnight. Next day, cell were treated using vehicle or the corresponding dose of cEt-ASO and allowed to proliferate for a further 72 hours. At endpoint, cells were fixed with 4% paraformaldehyde solution in PBS, permeabilised using 0.1% Triton X-100 in PBS and stained using DAPI. Plates were imaged using the OperaPhenix imaging system (Revvity) and cell counting was performed using Harmony high-content image analysis software (Revvity). Data are expressed as proliferation index, corresponding to the fold change of cell growth between the control cells and the cET-ASO-treated cells at endpoint for each condition across independent experiments.

### Confocal imaging and colocalisation

KPC cells were seeded in µ-Slide 8 Well high slides (Ibidi) using 3×10^3^ cells per well. Cells were allowed to attach and proliferate overnight. For endosomal localisation experiments, cells were previously transfected in 6-cm plates using the corresponding plasmids and Lipofectamine2000 (ThermoFisher), and 24 hours post-transfection cells were seeded into the chambered slides. Next day, cells were treated with either vehicle (PBS) or cET-ASOs accordingly and allowed to proliferate overnight (16 hours). Next day, samples were fixed using a 4% paraformaldehyde solution in PBS, permeabilised with 0.1% Triton X-100 solution in PBS. Next, samples were incubated with blocking buffer (5% normal goat serum in TBS) for 1 hour at RT. Primary antibodies were incubated overnight at 4°C diluted in blocking buffer. Next day, samples were washed three times in TBS buffer and further incubated with the corresponding secondary antibodies and DAPI diluted in blocking buffer for 1 hour at RT. Samples were next washed using TBS three times and soft mounting medium (Vectashield) was added to each well. Samples were visualised using either a Zeiss LSM880 microscope with Airyscan or a Zeiss Elyra 7 lattice SIM microscope, using a 63x/1.4NA objective and Zen Black software (ver2.3 for Airyscan, 3.0 for Elyra). For Airyscan imaging, frame-sequential capture was used and channel-specific bandpass filters as follows: blue: 420-480 + LP605; green: 420-480 + 495-550; red: 420-480 + 495-620; far-red: 570-620 + LP645. Pixel size was 0.04 µm. Airyscan 3D processing was used at default sharpness. For SIM, lattices used were: for AlexaFluor488 G5 (27.5µm); AlexaFluor568 G4 (32µm) or G5 (27.5µm) if phase modulation was judged to be sufficiently good; AlexaFluor647 G3 (36.5µm), for DAPI G6 (23µm) with 13 phases. z-stacks were captured at intervals of 91-125nm depending on the experiment; or for some samples at 55nm (over-sampling). Lasers (488nm, 561nm and 642nm, all 500mW; 405nm 50mW) were typically used at 2.5-4% power. Emitted light was captured on PCO edge 4.2M sCMOS cameras via a duolink adaptor, using a dual band-pass filter (490-560 + LP640) with a 50ms exposure time. Processing was by SIM^2^ using “standard live” settings of input SNR medium, iterations 16, regularisation 0.065, input and output sampling was set to x4, median filter. Grating period was 718.33nm and the resulting xy scaling was 0.031µm. Image analysis was performed using Fiji-ImageJ software to obtain Pearson correlation coefficients, and fluorescence intensity profiles of representative vesicle examples. Fluorescence intensity profiles are shown as the fold change relative to the maximum intensity value for each fluorophore. For 3D image reconstruction, Imaris software was used to create the 3D surfaces using the SIM^2^ data generated after processing the images obtained using the Elyra7 microscope.

### siRNA knockdown of scavenger receptors

H1299 cells were transfected with scavenger receptor siRNA pools using nucleofection Kit V and an AMAXA nucleofector (Lonza). 1.5 x 10^5^ cells were then seeded onto 6cm dishes and allowed to attach overnight. Next day, cells were incubated for 72h with the corresponding doses of cEt-ASO. Cells were then lysed and either protein or RNA was harvested. Effectivity of siRNA knockdown as well as *KRAS* mRNA targeting and downstream signalling by cEt-ASOs were assessed by either Western blotting or real-time PCR following the protocols described above.

### CRISPR knockout

Suppression of specific gene expression and generation of stable cell lines was achieved using the CRISPR-knockout genome editing system. Cloning of guide RNAs into lentiviral plasmids and generation of stable cell lines after lentiviral infection was performed accordingly to the protocol established by the Zhang lab^64^. Briefly, guide RNAs were designed, annealed and inserted into lentiCRISPRv2 lentiviral plasmids (Addgene #52961). Lentiviral particles were produced by transducing HEK293FT cells with the lentiCRISPRv2 guideRNA-expressing plasmid together with psPAX2 and VSV-G packaging plasmids. Supernatants were collected after 48 hours and transferred onto the recipient cells. Selection of positively transduced cells was assessed with antibiotic selection (puromycin) in the culture media. After three passages, the successful knock out of the target genes was assessed by qPCR as well as by Western blotting.

### EPHA2-TurboID pulldown and immunofluorescence

KPC cells stably expressing an EPHA2-TurboID construct^13^ were seeded in 10cm plates using 2×10^6^ cells per plate. Cells were allowed to attach and proliferate overnight. Next day, cells were incubated with 100µM biotin and 5µM cET-ASO for 16 hours. Next day, cells were placed on ice to stop the biotin ligation reaction and washed five times with ice-cold PBS. Cells were scrapped from the plate in PBS and pelleted at 300x*g* for 3 minutes at 4°C. Cells were lysed in RIPA buffer supplemented with protease inhibitors and sonicated. The insoluble fraction was discarded and the free biotin excess from the lysates was removed using an AMICON 3K filter column (Millipore). Cell lysates were recovered and further incubated with streptavidin-agarose beads (Millipore) for 2 hours at 4°C. After the incubation, the beads were further washed sequentially with RIPA, 1M KCl, 0.1M Na2CO3, 2M urea and a final 100mM NaCl - 10mM Tris pH7.5 – 0.5mM EDTA buffer. Beads were then resuspended in loading buffer and 2mM biotin and biotinylated targets were assessed using western blotting.

For immunofluorescence, KPC cells stably expressing an EPHA2-TurboID construct were plated onto glass bottom 35mm dishes and allowed to attach overnight. Next day, cells were treated with biotin as described above and 16h afterwards, biotin was washed five times with PBS and the cells were fixed using 4% paraformaldehyde. Cells were processed as described above and biotinylated proteins were assessed using streptavidin conjugated to Alexa488 (ThermoFisher). Samples were visualised in an Zeiss880 Airyscan microscope as described above.

### Genomic data

Publicly available data, analysis and graphs were obtained from cBioPortal^65^ using the TCGA dataset corresponding to Pancreatic adenocarcinoma (PAAD). GEPIA2 portal^66^ was used to obtain the comparison between tumour and normal tissue data. Further genomic data regarding mRNA expression in PDAC was obtained from analysis of microarray and RNAseq data from Bailey et al.^33^, as well as PDAC subtype classification expression data.

## Acknowledgements

We would like to thank core staff in the Biological Services Unit, the Beatson Advanced Imaging Resource (BAIR), Molecular Services, Histology Facility and Central Services (CRUK Scotland Institute) for their support which facilitated the work described in this manuscript. The results shown here are in whole or part based upon data generated by the TCGA Research Network: https://www.cancer.gov/tcga. The data used for the analyses described in this manuscript were obtained from the [GTEx Portal](https://gtexportal.org/home/) and/or [dbGaP](https://www.ncbi.nlm.nih.gov/gap/) accession number phs000424.vN.pN. This paper was critically reviewed by Catherine Winchester (CRUK Scotland Institute).

## Author contributions

Experimental design: SM, JCN, PW & MB

Investigation: SM, LMcG, PT, DT, & SA

Funding acquisition: JCN & CB

Writing – original draft: SM & JCN

Writing – review & editing: All authors

## Data and materials availability

The data supporting the findings of this study are available within the article and its supplementary information files and from the corresponding author upon request.

## Supplementary figure legends

**Figure S1 (related to Figure 1).**
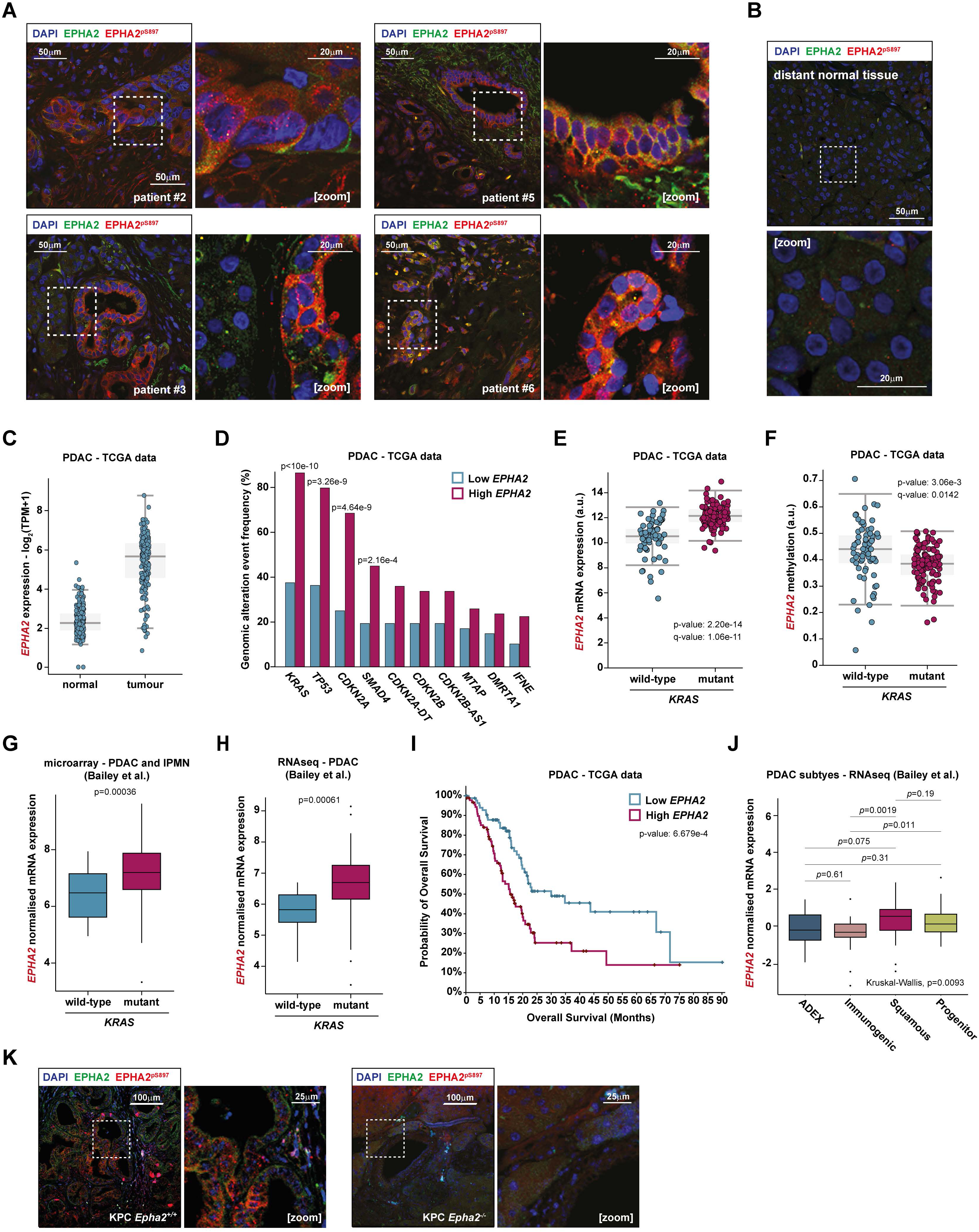
(A) PDAC patient tumour tissue immunostaining. Left panels show staining of EPHA2 (green) and Ser^897^ phosphorylated EPHA2 (red) in transformed cells. Right panels correspond to the amplification of the area inside the white dotted line frame, showing the detailed localisation of the staining. (B) Top panels shows the absence of both EPHA2 and Ser^897^ phosphorylated EPHA2 in non-transformed adjacent tissue in human pancreas. Bottom panel corresponds to amplification of the area inside the white dotted line frame. (C) *EPHA2* expression in normal pancreas (TCGA and GTEx, n=171) compared to expression in PDAC (TCGA, n=179). (D) Analysis of the mutational frequency in the most common oncodrivers in PDAC (TCGA), comparing low *EPHA2* expression tumours (blue, n=88) with high *EPHA2* expression tumours (magenta, n=89). Two-sided Fisher exact test. (E) Comparison of the abundance of *EPHA2* mRNA in either *KRAS* wild-type (blue, n=67) or *KRAS* mutant (magenta, n=117) PDAC tumours (TCGA). Students t-test. (F) Comparison of the promoter methylation levels on the *EPHA2* gene in either *KRAS* wild-type (blue, n=67) or *KRAS* mutant (magenta, n=117) PDAC tumours (TCGA). Students t-test. (G) Comparison of the abundance of *EPHA2* mRNA in either *KRAS* wild-type (blue) or *KRAS* mutant (magenta) PDAC tumours (microarray data, PDAC and intraductal papillary mucinous neoplasm – IPNM). *Kras* wild-type n=24, *Kras* mutant n=207. Students t-test. (H) Comparison of the abundance of *EPHA2* mRNA in either *KRAS* wild-type (blue) or *KRAS* mutant (magenta) PDAC tumours (RNAseq data^33^). *Kras* wild-type n=11, *Kras* mutant n=84. Students t-test. (I) Survival plot - Probability of overall survival of PDAC patients comparing patients with low *EPHA2* expression (blue, n=88) and patients with high *EPHA2* expression (magenta, n=89). Logrank Test. (J) *EPHA2* mRNA expression comparison across the different Bailey^33^ PDAC subtypes. ADEX n=16, Immunogenic n=25, Progenitor n=30, Squamous n=25. Kruskal-Wallis. (K) Tumour tissue obtained from wild-type KPC mice (*Kras^G12D/+^*, *LSL-Tp53^172H/+^*, *Pdx1-Cre*, left panels), and *Epha2* knockout KPC mice (*Kras^G12D/+^*, *LSL-Tp53^172H/+^*, *Pdx1-Cre, Epha2^-/-^*, right panels). *Epha2* knockout PDAC tumours show the absence of both EPHA2 and Ser^897^ phosphorylated EPHA2 staining. Image zooms correspond to the area inside the dotted box.

**Figure S2 (related to Figure 1).**
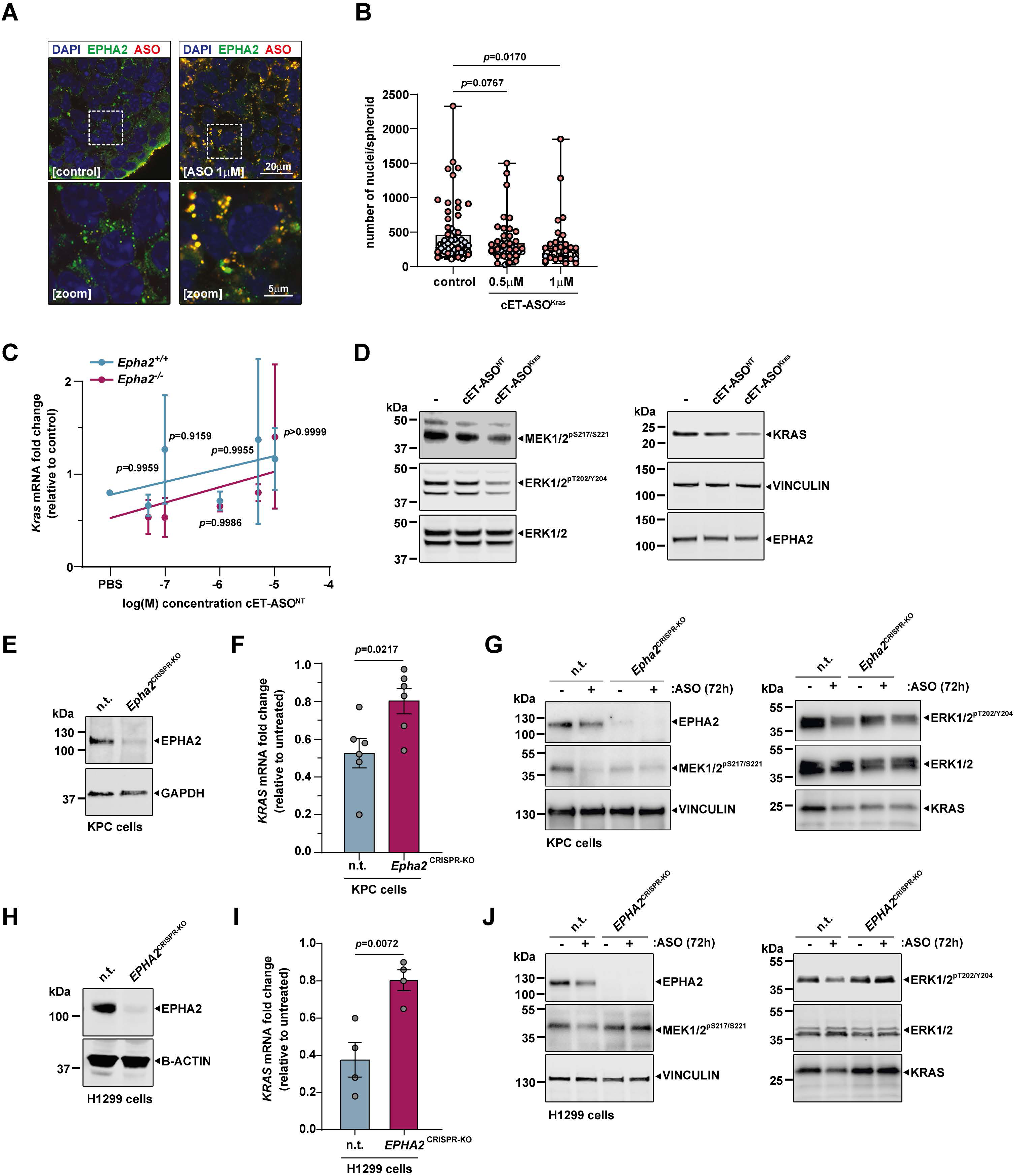
(A) High-resolution micrograph of KPC cells either untreated (control) or treated with cET-ASO^Kras^ for 72h. EPHA2 (green) and cET-ASO^Kras^ (red) staining are shown overlapping in intracellular vesicles. Bottom panels correspond to an image amplification of the area inside the white dotted line frames in the top panels. (B) High-content image analysis of the number of nuclei in each KPC spheroid. Dot colouring indicates each independent experiment (n=3). One-way ANOVA, Dunnett multiple comparison test. (C) Dose response curve of wild-type (*Epha2^+/+^,* blue) or *Epha2* knockout (*Epha2^-/-^*, magenta) KPC cells of *Kras* mRNA expression after 72 hours of treatment with cET-ASO^NT^. 2-way ANOVA, Sidak multiple comparison test, n=3 individual experiments. (D) Western blotting comparing the effect between cET-ASO^NT^ (nt) and cET-ASO^Kras^ (Kras) on MEK1/2 and ERK1/2 phosphorylation and KRAS expression in KPC cells. Vinculin was used as loading control. (E) Western blotting of KPC cell lysates obtained from either KPC cells transduced with a non-targeting sgRNA (nt) or a *Epha2*-targeting sgRNA (Epha2^CRISPR-KO^), showing the depletion of EPHA2 expression in the latter. GAPDH was used as loading control. (F) *Kras* mRNA expression in either non-targeting or Epha2^CRISPR-KO^ KPC cells after treatment with 0.5μM cET-ASO^Kras^ for 72h. Student t-test, n=5. (G) Western blotting in either non-targeting or Epha2^CRISPR-KO^ KPC cells after treatment with 0.5μM cET-ASO^Kras^ for 72h, showing the effect of the ASO on MEK1/2 and ERK1/2 phosphorylation and KRAS expression. Vinculin was used as loading control. (H) Western blotting of H1299 (non-small cell lung carcinoma) cell lysates obtained from either H1299 cells transduced with either a non-targeting sgRNA (nt) or a *EPHA2*-targeting sgRNA (EPHA2^CRISPR-KO^), showing the depletion of EPHA2 expression in the latter. Beta-actin was used as loading control. (I) *KRAS* mRNA expression in either non-targeting or EPHA2^CRISPR-KO^ H1299 cells after treatment with 0.5μM cET-ASO^Kras^ for 72h. Student t-test, n=4. (J) Western blotting in either non-targeting or EPHA2^CRISPR-KO^ H1299 cells after treatment with 0.5μM cET-ASO^Kras^ for 72h, showing the effect of the ASO on MEK1/2 and ERK1/2 phosphorylation and KRAS expression. Vinculin was used as loading control.

**Figure S3 (related to Figure 3).**
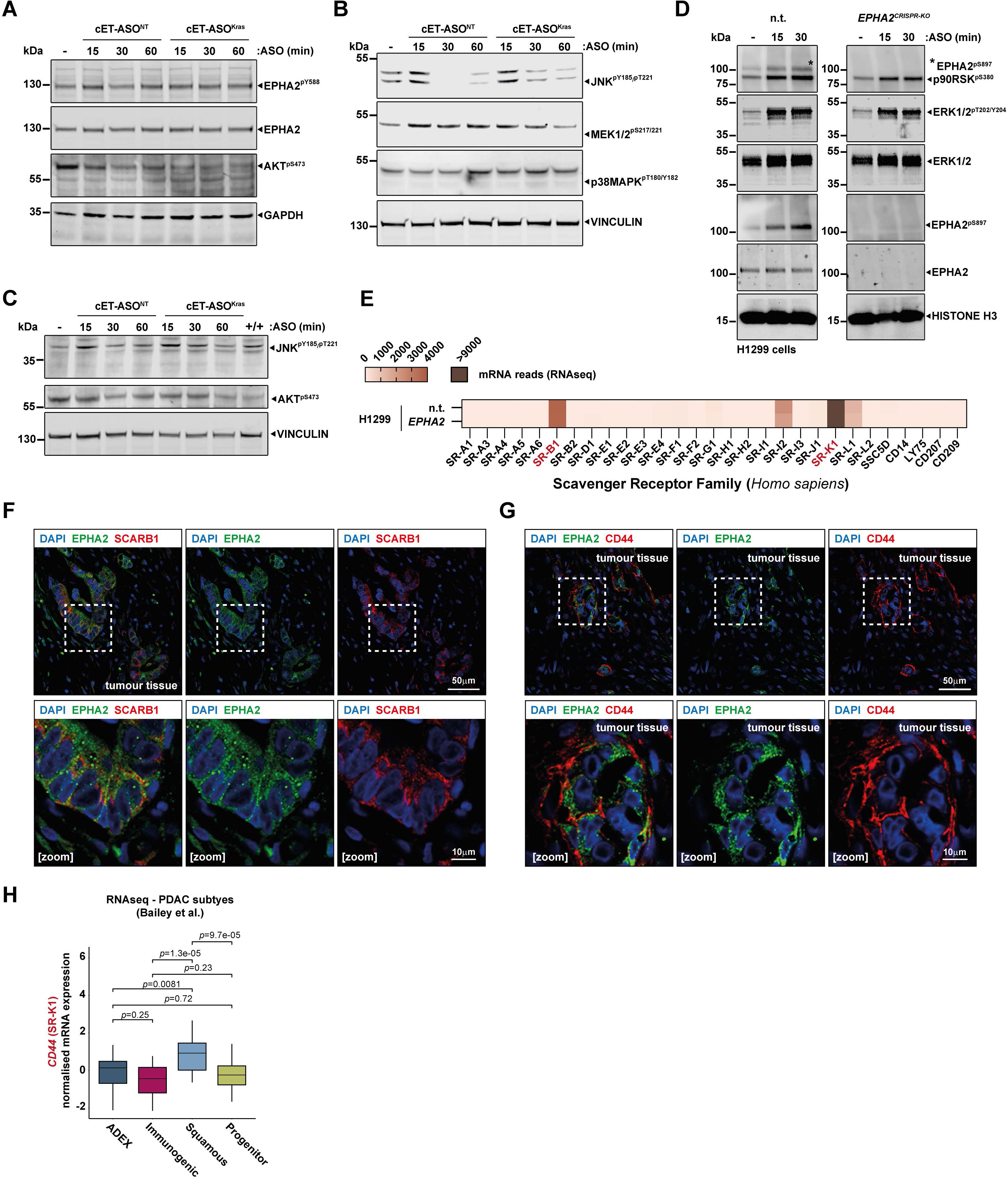
(A) Western blotting of either non-targeting cET-ASO^NT^ or cET-ASO^Kras^-treated KPC cells, showing the absence of phosphorylation induction on EPHA2 Tyr^588^ and AKT Ser^473^. GAPDH was used as loading control. (B) Western blotting of either non-targeting cET-ASO^NT^ or cET-ASO^Kras^-treated KPC cells, showing the induction of phosphorylation on JNK Tyr^185^ and Thr^221^, MEK1/2 Ser^217^ and Ser^221^, as well as the lack of induction of phosphorylation on p38MAPK Thr^180^ and Tyr^182^. Vinculin was used as loading control. (C) Western blotting in either non-targeting cET-ASO^NT^ or cET-ASO^Kras^-treated KPC cells, showing the induction of phosphorylation on JNK Tyr^185^ and Thr^221^ and the lack of induction of phosphorylation on AKT Ser^473^ in both KPC *Epha2*^+/+^ and *Epha2*^-/-^ cells. These panels correspond to the same western blot as in Fig.3D, including vinculin as a loading control. Histone H3 was used as loading control. (D) Western blotting showing the induction of the phosphorylation on EPHA2 Ser^897^, p90RSK Ser^380^, and ERK1/2 Tyr^202/204^ in cET-ASO^Kras^-treated *EPHA2*^+/+^ and *EPHA2*^CRISPR-KO^ H1299 cells. * Band corresponds to the band previously obtained by blotting for EPHA2 phospho Ser^897^. (E) Heatmap showing the levels of mRNA expression (RNAseq normalised reads) of each member of the scavenger receptor family in either *EPHA2^+/+^* (top) or *EPHA2-* siRNA depleted (bottom) H1299 cells. Red-labelled names correspond to the highest expressed genes both in KPC and H1299 cells. Outlier values are indicated using dark brown colour. (F) PDAC patient tumour tissue immunostaining. Top panels show the staining of EPHA2 (green) and SR-B1 (SCARB1, red) within the tumour nodules. Bottom panels correspond to the amplification of the area contained in the white dotted line frame. (G) PDAC patient tumour tissue immunostaining. Top panels show the staining of EPHA2 (green) and SR-K1 (CD44, red) within the tumour nodules. Bottom panels correspond to the amplification of the area contained in the white dotted line frame. (H) *CD44* mRNA expression comparison across the different Bailey PDAC subtypes. Kruskal-Wallis.

**Figure S4 (related to Figure 3).**
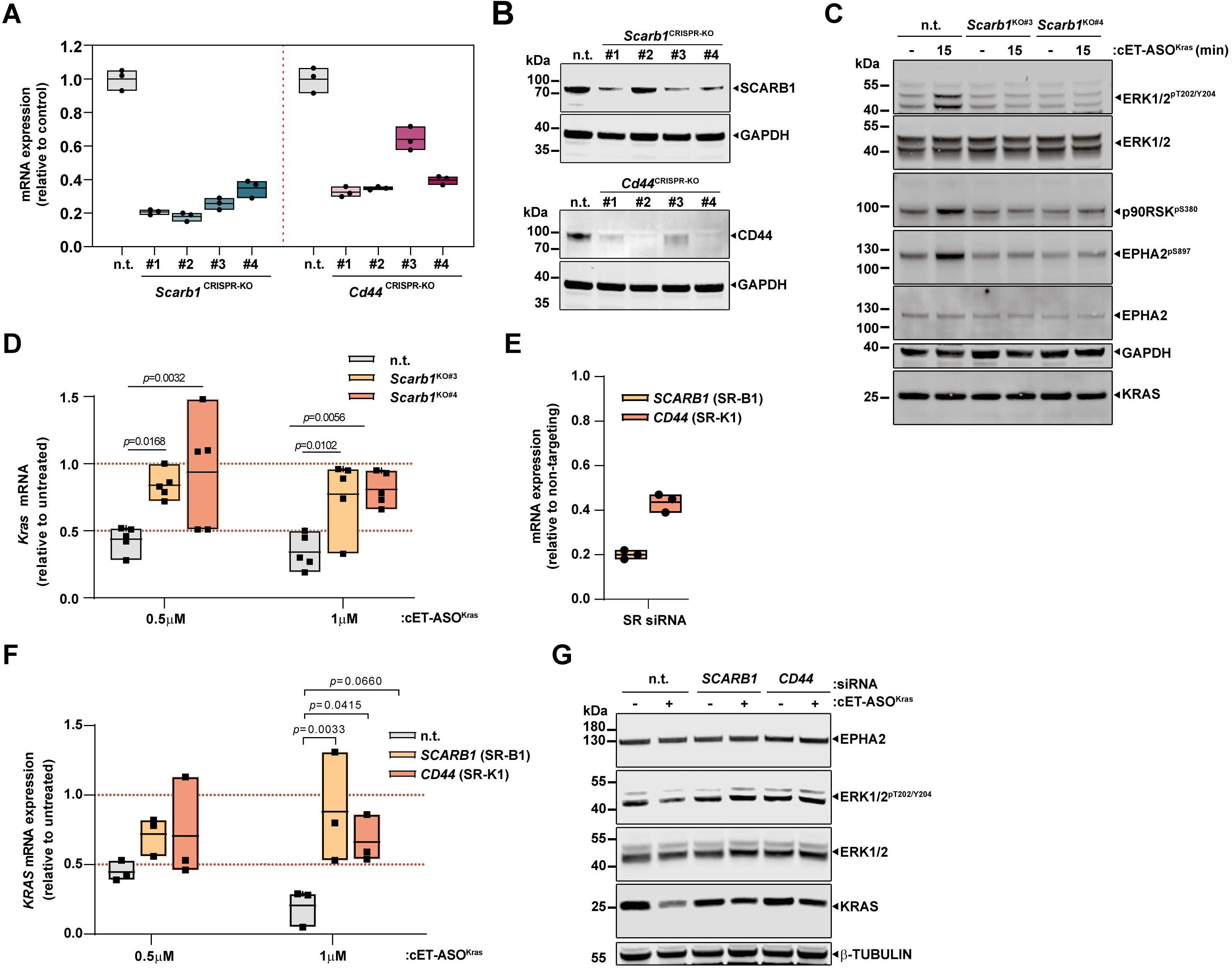
(A) *Scarb1* and *Cd44* mRNA expression levels in KPC cells transduced with either a non-targeting sgRNA (n.t.) or four different sgRNAs targeting either *Scarb1* (*Scarb1*^CRISPR-KO^) or *Cd44* (*Cd44*^CRISPR-KO^). Dots represent n=3 independent experiments. (B) Western blotting showing the protein expression of either SCARB1 in non-targeting and *Scarb1*^CRISPR-KO^ KPC cells, or CD44 in *Cd44*^CRISPR-KO^ KPC cells. GAPDH was used as loading control. (C) Western blotting showing the effect of *Scarb1* depletion in *Scarb1*^CRISPR-KO^ KPC cell lines on the cET-ASO^Kras^-induced activation of ERK1/2 Tyr^202/204^, p90RSK Ser^380^ and EPHA2 Ser^897^ phosphorylations. GAPDH was used as loading control. (D) *Kras* mRNA expression in either non-targeting or *Scarb1^CRISPR-KO^* KPC cell lines treated with either 0.5μM or 1μM cET-ASO^Kras^ for 72h. 2-way ANOVA, Dunnett multiple comparison test, n=5. (E) *SCARB1* and *CD44* mRNA expression levels in H1299 cells transduced with either a non-targeting siRNA or a pooled siRNA for each respective scavenging receptor. Results expressed as the fold change of each mRNA in non-targeting transduced cells. (F) *KRAS* mRNA expression in H1299 cells transduced with either a non-targeting siRNA or a pooled siRNA for each respective scavenging receptor, treated with either 0.5μM or 1μM cET-ASO^Kras^ for 72h. 2-way ANOVA, Tukey multiple comparison test, n=3. (G) Western blotting showing the effect of depletion of *SCARB1* and *CD44* scavenger receptors in H1299 cell lines, on the MAPK signalling activation and *KRAS* expression after 72h of cET-ASO^Kras^ addition. Beta-tubulin was used as loading control.

**Figure S5 (related to Figure 4).**
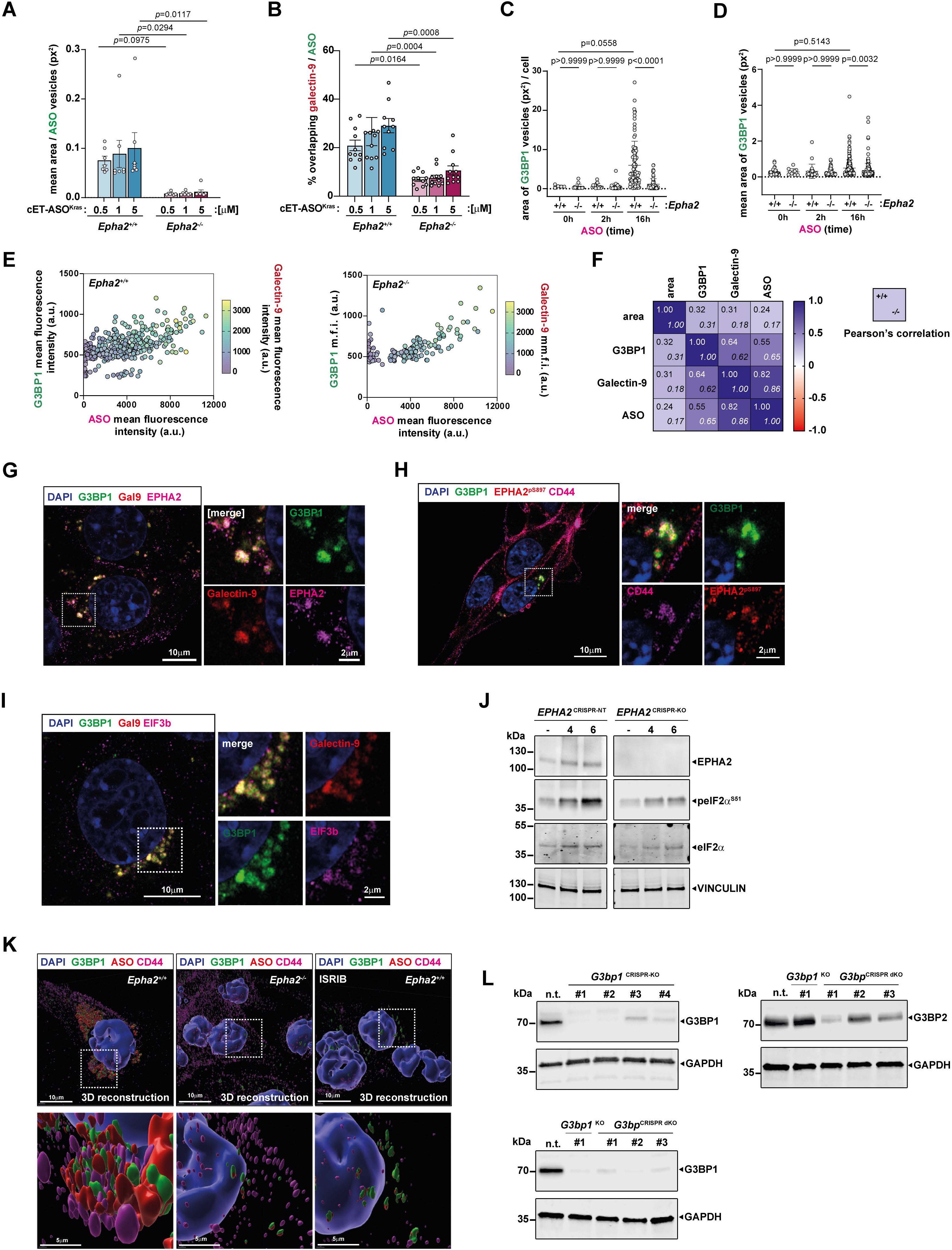
(A) Quantification of the mean area size of ASO-positive vesicles in either *Epha2^+/+^*(blue) or *Epha2^-/-^* (magenta) KPC cells, after treatment with either 0.5, 1 or 5 μM concentrations of cET-ASO^Kras^ for 16 hours. One-way ANOVA, Tukey multiple comparison test, n=7. (B) Quantification of the mean area size of galectin-9-positive vesicles in either *Epha2^+/+^* (blue) or *Epha2^-/-^* (magenta) KPC cells, after treatment with either 0.5, 1 or 5 μM concentrations of cET-ASO^Kras^ for 16 hours. One-way ANOVA, Tukey multiple comparison test, n=11. (C) Average sum of the area of G3BP1-positive structures per cell, in either *Epha2^+/+^* or *Epha2^-/-^* KPC cells after incubation with cET-ASO^Kras^ for 0, 2 or 16 hours., and (D) mean area of G3BP1-positive structures per cell, in either *Epha2^+/+^* or *Epha2^-/-^* KPC cells after incubation with cET-ASO^Kras^ for 0, 2 or 16 hours. Dots correspond to different fields of view obtained in three individual experiments. One-way Anova – Tukey test. (E) Scatter plot of the mean intensity values for G3BP1, ASO and galectin-9 for each cell in either *Epha2^+/+^* (left panel) or *Epha2^-/-^* (right panel) KPC cells after incubation with cET-ASO^Kras^ for 16 hours. (F) Pearson’s correlation coefficient heatmap of the mean intensity values for G3BP1, ASO and galectin-9 for each cell in either *Epha2^+/+^* (top value) or *Epha2^-/-^* (bottom) KPC cells after incubation with cET-ASO^Kras^ for 16 hours. (G) Micrographs of immunofluorescence confocal imaging showing the overlap between G3BP1-GFP, EPHA2 and galectin-9 in intracellular structures in KPC cells treated with cET-ASO^Kras^ for 16h. (H) Micrographs of immunofluorescence confocal imaging showing the overlap between G3BP1-GFP, internalised EPHA2^pS897^ and intracellular CD44 in KPC cells treated with cET-ASO^Kras^ for 16h. Image zooms correspond to the area inside the corresponding dotted line boxes. (I) Micrographs of immunofluorescence confocal imaging showing the overlap between G3BP1-GFP, galectin-9 and eIF3b in intracellular structures in KPC cells treated with cET-ASO^Kras^ for 16h. (J) Western blotting in of either non-targeting or EPHA2^CRISPR-KO^ H1299 cells showing the effect on eIF2a phosphorylation after incubation with ASO cET-ASO^Kras^ for 4 or 6 hours. Vinculin was used as loading control. (K) 3D reconstruction of immunofluorescence confocal imaging showing the overlap between G3BP1-GFP, internalised cET-ASO^Kras^ and intracellular CD44 in either *Epha2^+/+^*, *Epha2^-/-^* or ISRIB (1μM) treated *Epha2^+/+^* KPC cells. (L) Western blotting showing the protein expression of G3BP1 in non-targeting (n.t.) and KPC cells transduced with four different sgRNA targeting *G3bp1* in KPC cells (*G3bp1*^CRISPR-KO^, upper left panel); western blotting showing the protein expression of G3BP2 in non-targeting (n.t.), *G3bp1*^CRISPR-KO^ ^#1^ and *G3bp1*^CRISPR-KO^ ^#1^ KPC cells transduced with three different sgRNA targeting *G3bp2* (*G3bp1/2*^CRISPR-KO^, upper right panel); western blotting showing the protein expression of G3BP1 in non-targeting (n.t.), *G3bp1*^CRISPR-KO^ ^#1^ and *G3bp1*^CRISPR-KO^ ^#1^ KPC cells transduced with three different sgRNA targeting *G3bp2* (*G3bp1/2*^CRISPR-KO^, lower left panel). GAPDH was used as loading control.

**Figure S6 (related to Figure 2 and discussion).**
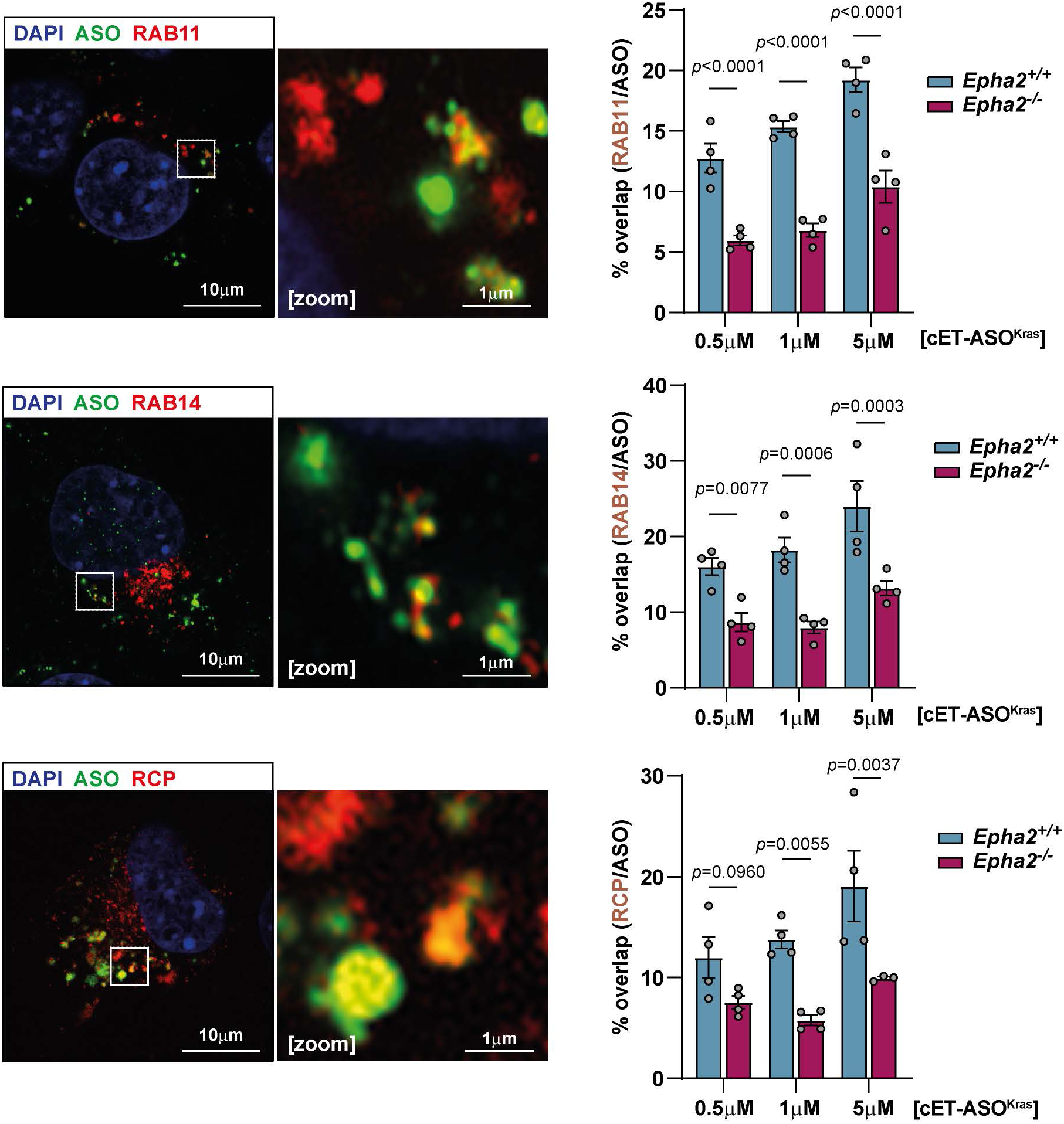
(A) Immunofluorescence confocal imaging of the overlap between cET-ASO^Kras^ (green) and Rab11-positive vesicles (top panel, red), Rab14-positive vesicles (middle panel, red) or Rab-coupling protein (RCP) (bottom panel, red), inside KPC cells after 16 hours of incubation (left panel). Image amplification of the selected area (dotted line frame), and quantification of cET-ASO^Kras^ and the different endosomal markers in either KPC *Epha2^+/+^* (blue) or *Epha2^-/-^*(magenta) cells treated at different ASO concentrations for 16h (corresponding right panels). 2-way ANOVA, n=4.

## Key resources table

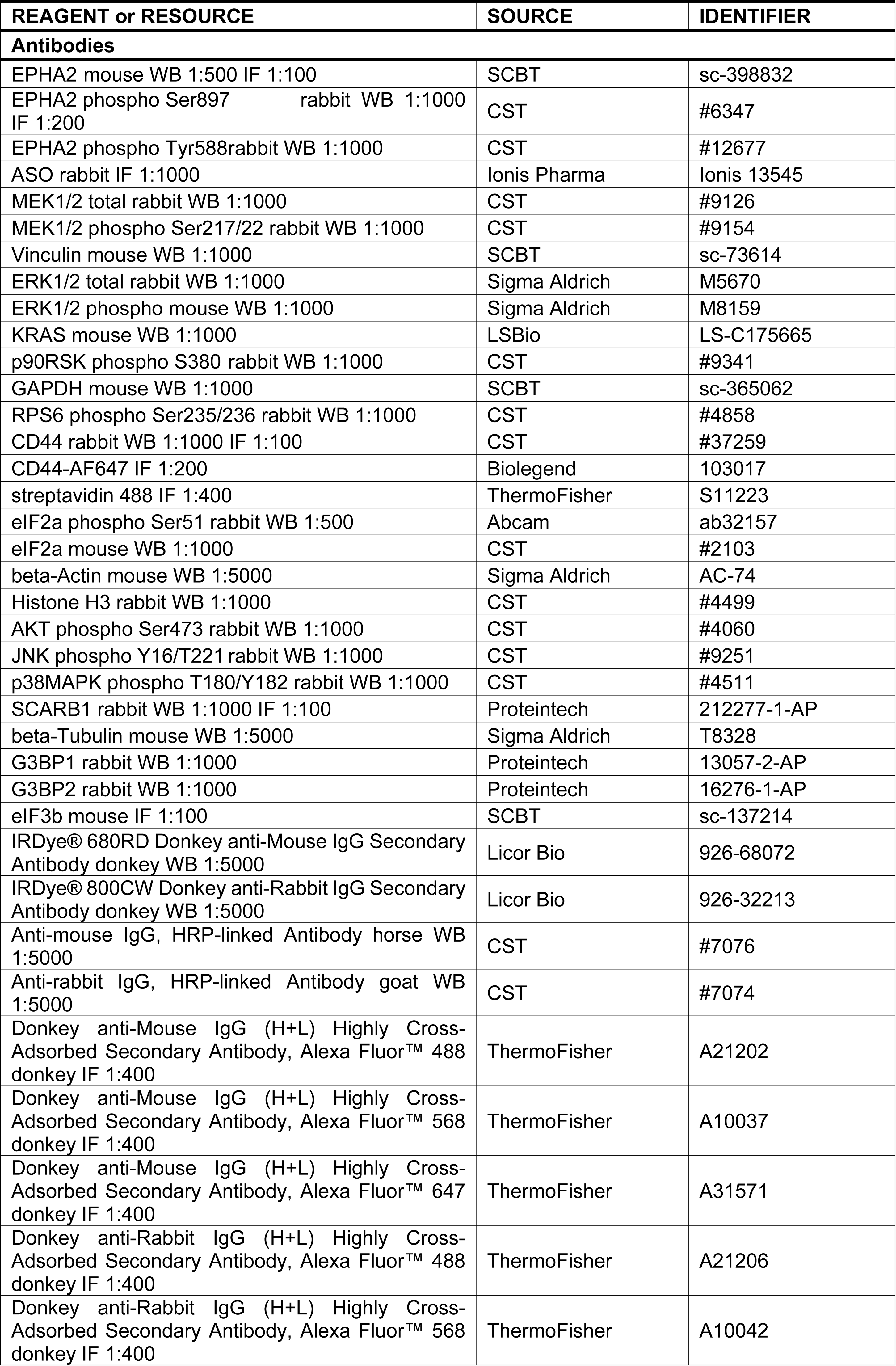
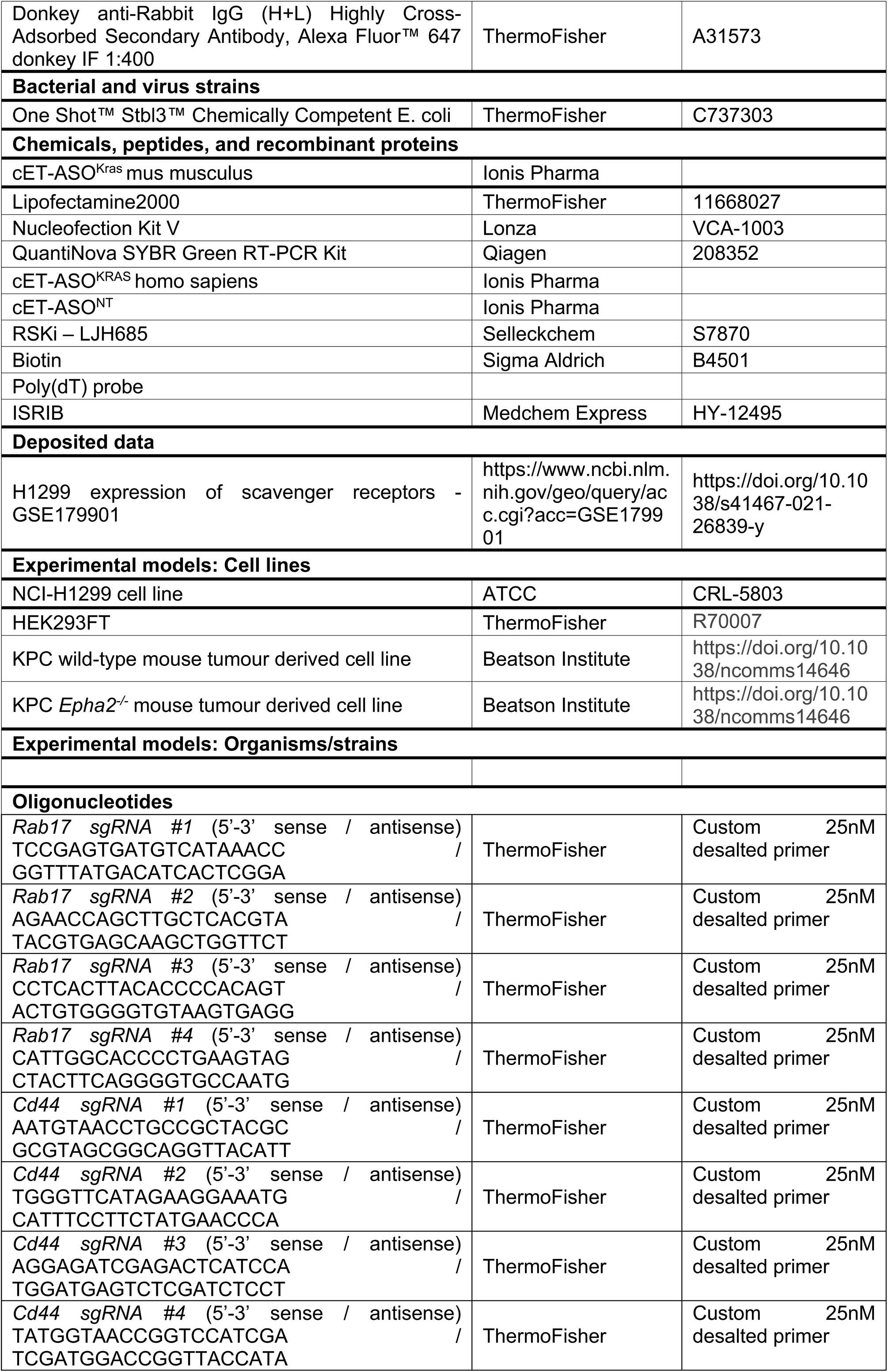
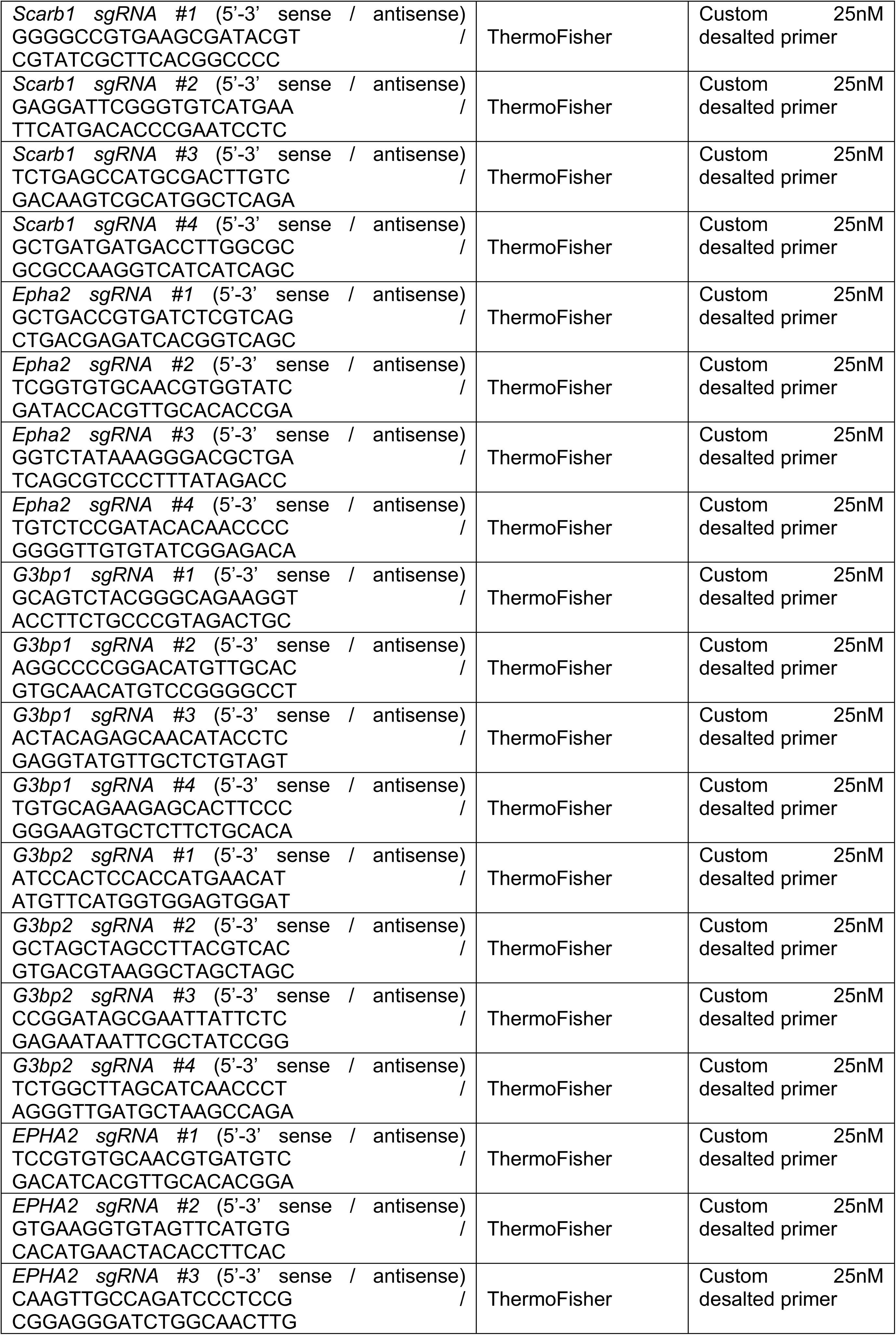
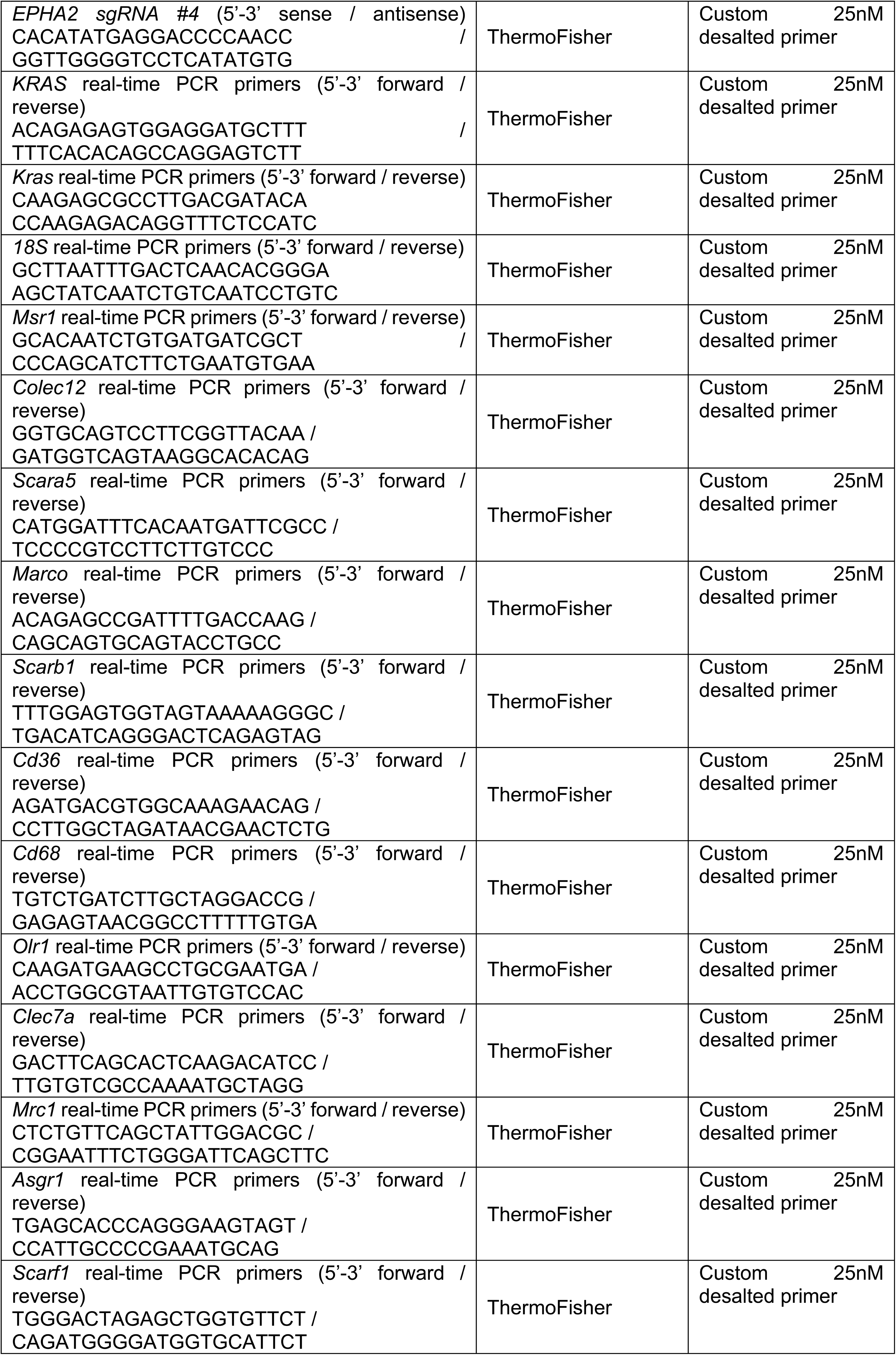
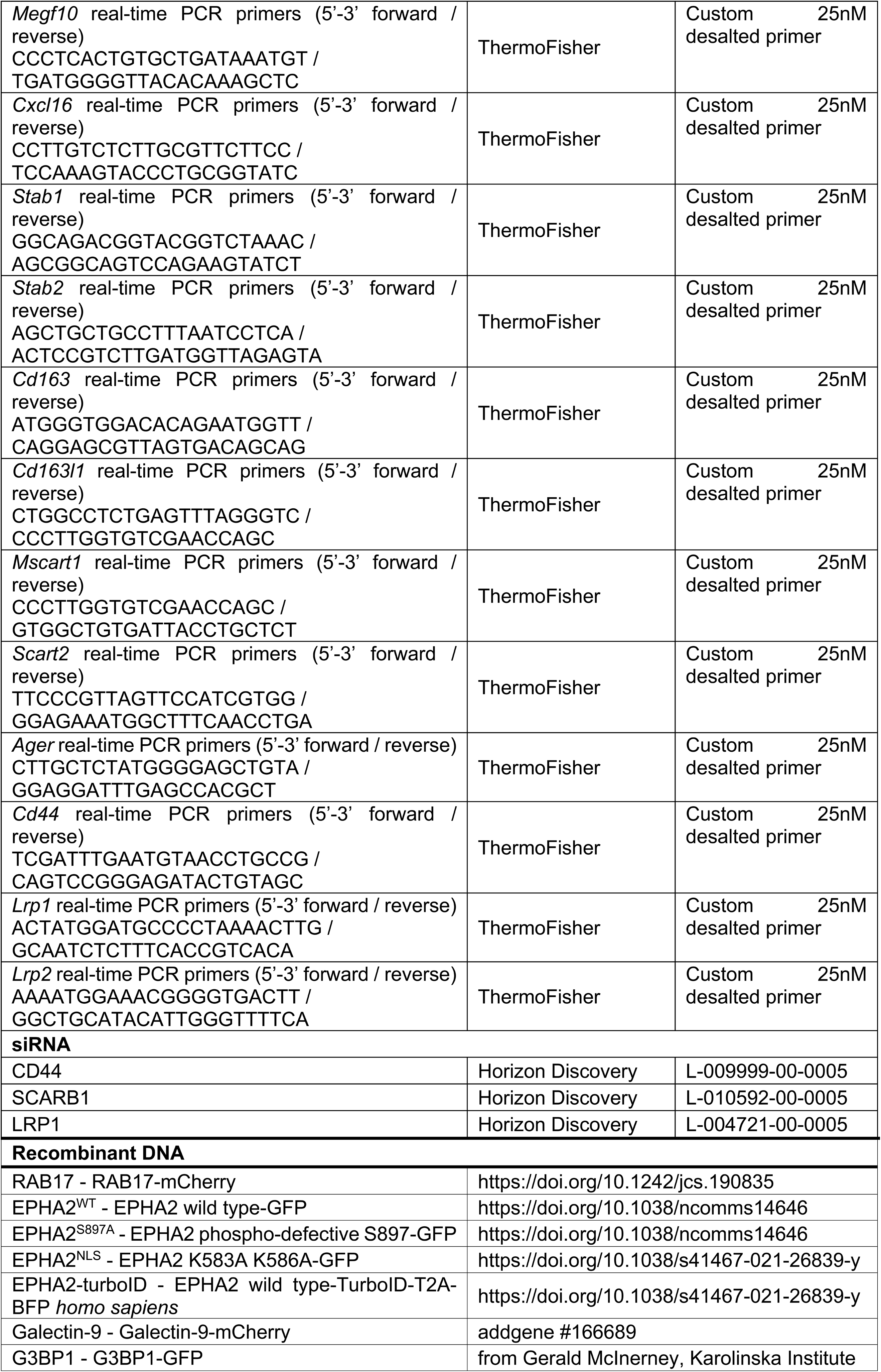
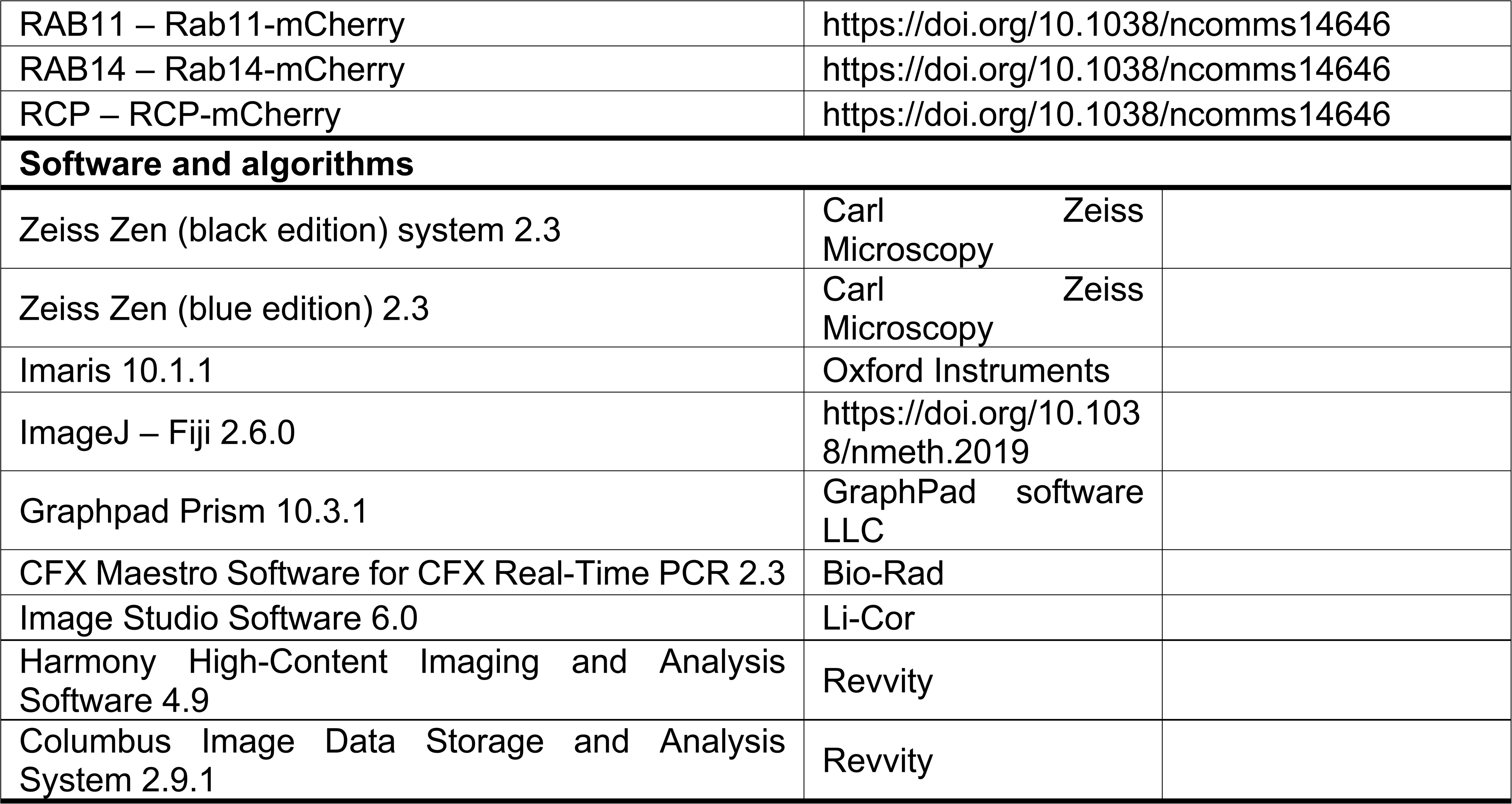

